# Compartment coupling integrates patterning and morphogenetic information during development

**DOI:** 10.1101/2024.09.16.613243

**Authors:** Anubhav Prakash, Sukanya Raman, Raman Kaushik, Pallavi Manchanda, Anton S Iyer, Raj K. Ladher

**Affiliations:** National Centre for Biological Sciences, Tata Institute of Fundamental Research, GKVK PO, Bellary Road, Bangalore, India, 560065; Trivedi School of Biosciences, Ashoka University, Plot No. 2, Rajiv Gandhi Education City, National Capital Region P.O. Rai, Sonipat Haryana-131029

**Author notes:** Authors for Correspondence: **Anubhav Prakash**, **Raj K. Ladher**.

**Keywords:** Planar Cell Polarity, Adhesion-code, Inner Ear, Compartment-coupling, Vinculin

## Abstract

Morphogenetic information arises from a combination of genetically encoded cellular properties and emergent cellular behaviours. The spatio-temporal implementation of this information is critical to ensure robust, reproducible tissue shapes, yet the principles underlying its organisation remain unknown. We investigated this principle using the mouse auditory epithelium, the organ of Corti (OC). OC consists of a sensory domain, which transduces sound through polar mechanosensory hair cells (HC), part of a mosaic with supporting cells (SC). On either side of the sensory domain are non-sensory domains. These domains undergo cellular rearrangements, which, together, lead to a spiral cochlea that contains planar polarised HCs. This makes the mammalian cochlea a compelling system to understand coordination across spatial scales. Using genetic and ex-vivo approaches, we found patterning of OC into sensory and non-sensory domains is associated with a combinatorial expression of adhesion molecules, which underpins OC into spatially defined compartments, enabling planar cell polarity (PCP) cues to regulate compartment-specific organisation. Through compartment-specific knockouts of the PCP protein, Vangl2, we find evidence of compartment coupling, a non-linear influence on the organisation within one compartment when cellular organisation is disrupted in another. In the OC, compartment coupling originates from vinculin-dependent junctional mechanics, coordinating cellular dynamics across spatial scales.

**Graphical Abstract:** 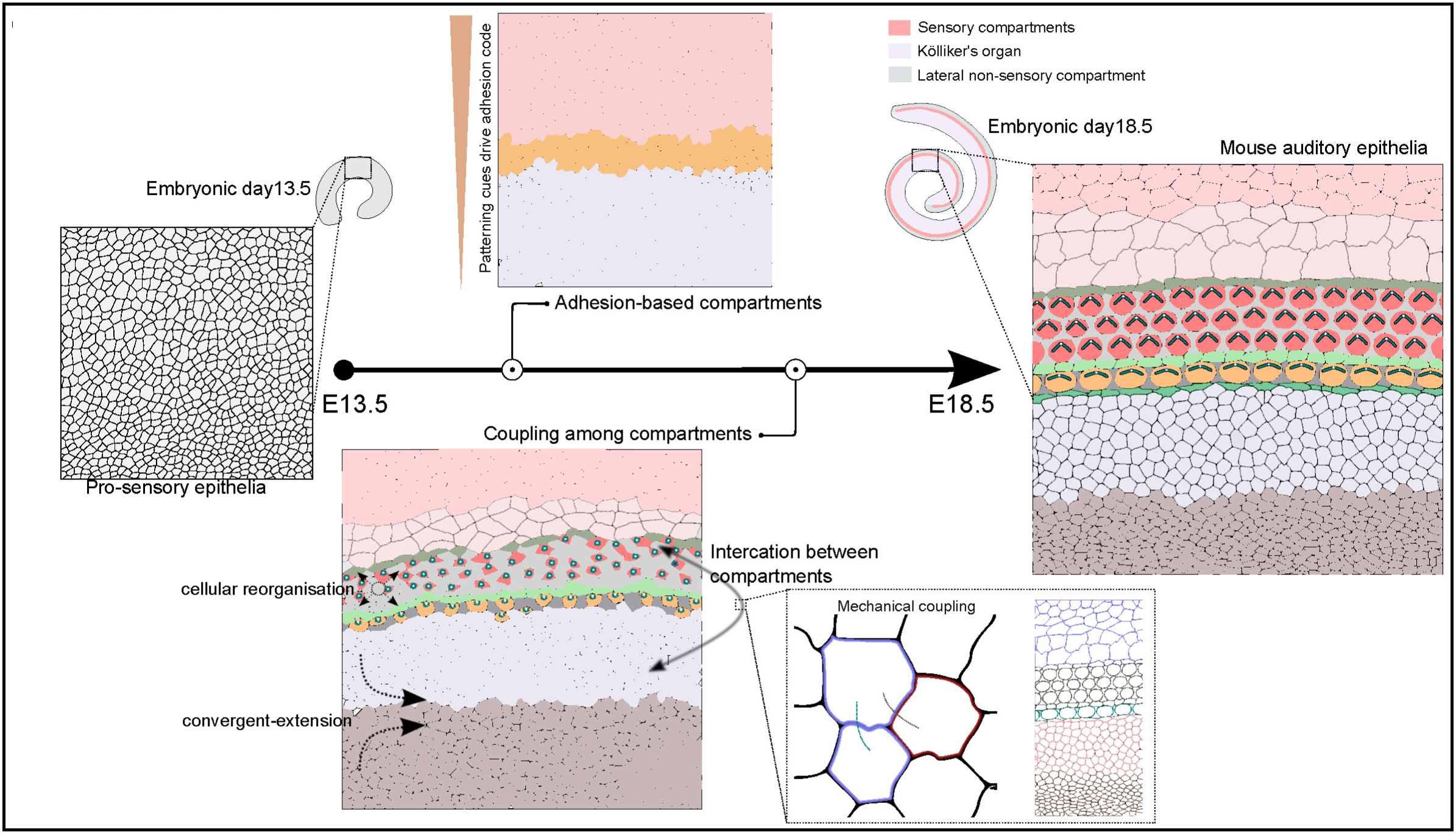

## INTRODUCTION

The reproducibility and robustness of morphogenesis result from carefully implemented developmental instructions. Through a combination of signalling cues, gene networks, and self-organising mechanisms, patterning divides an organogenic field into compartments [1, 2]. While compartments allow the segregation of distinct cell behaviors, these behaviors must be spatiotemporally coordinated across compartments so that functional organs, with the correct shape and pattern, forms. The mouse auditory epithelia is an excellent system to investigate this unknown coordination.

The mouse auditory epithelia, called the organ of Corti (OC) is found within the cochlea. It is a spiral-shaped organ and is responsible for detecting and transducing sound across a wide spectrum of frequencies [3, 4]. Sound is transduced by hair cells (HC) through an asymmetric hearing organelle, the hair bundle on its apical surface. HCs are of two types, inner HCs (IHC) arranged into a single row that transmits information to brain; and outer HCs (OHCs) arranged into three rows that amplifies the mechanical input. Both these HCs are intercalated by supporting cells (SCs) and together forms the sensory domain of OC. This sensory domain is flanked by non-sensory compartments. On the medial side (inner edge of spiral) is Kölliker’s organ (KO) and on the lateral side (outer edge of spiral) is a lateral non-sensory compartment, which includes Hensen’s and Claudius’ cells (Fig. S1A).

The cochlea is initially apparent as a ventral out-pocketing of the otocyst. Radial patterning of the nascent cochlear duct by morphogen signalling establishes non-sensory and sensory domains of the OC [5, 6]. The cells in the sensory domain become post-mitotic and, through juxtracrine signalling, differentiate into a mosaic of HC and SC [7, 8]. As HC develop, they form asymmetrically localised hair bundles which align with the tissue axis, a process known as planar polarity. Both experimental approaches and mathematical modelling have shown that local coordination among HCs and SCs driven by differential junctional tension could drive the organisation of HCs and align HC polarity to the tissue axis [9-11, 12. While cells locally coordinate in the sensory domain, large-scale convergence and extension (CE) movements together with growth and proliferation cause the OC to elongate contributing to morphogenesis and the spiralling of the OC{Yamamoto, 2009 #2047, 13-17]. Thus, local domain-specific processes that order HC must integrate with large-scale tissue remodelling. How they integrate is not known.

The OC compartmentalises into smaller domains that show a combinatorial expression of Cadherin 1, 2, and 4 [14, 18, 19]. Using mutants of fibroblast growth factor signalling and ex-vivo cultures, we find that the adhesion code ensures compartment integrity during convergent and extension movements. Each compartment uses the PCP molecule, Vangl2 to develop a distinct cellular organisation. Using compartment-specific knockouts of Vangl2, we find that cellular organisation within each compartment has a non-linear influence on the organisation of another compartment, a novel phenomenon called compartment coupling. In mice mutant for the junctional force transmission component Vinculin, we show that compartment coupling has a mechanical component. Our work suggests that compartment coupling underpins the integration of cellular ordering with large-scale tissue remodelling. Given the widespread use of compartments, inter-compartment coupling is likely a fundamental feature of the morphogenesis of many developing tissues and organs.

## Results

### Domain organisation is preserved during cochlear elongation

To investigate the mechanism that could integrate local organisation with tissue scale remodeling, we first asked how organisation in OC evolves from embryonic day (E)15.5, when the HCs are first apparent to E18.5, when the HCs achieve their final organisation. Immunostaining for marker of HCs, Myosin 7a, shows the sensory domain is already established by E15.5 (Fig. S1B). Positive immunostaining for p75NTR, a molecular marker for inner pillar cells which segregates the IHCs from OHCs shows the presence of medial and lateral sensory domains at this stage (Fig. S1C). Further, the expression of brain lipid binding-protein (BLBP) a molecular marker for Hensen’s cells (HnC, a SC type lateral to OHC) and Inner phalangeal cells (IPhC, SC type intercalating IHC) showed the sensory and the non-sensory domains are established by E15.5 (Fig. S1D). This organisation of OC into medial and lateral non-sensory and sensory domains suggests that by E15.5 the OC is radially patterned (Fig. 1B).

**Figure 1.**
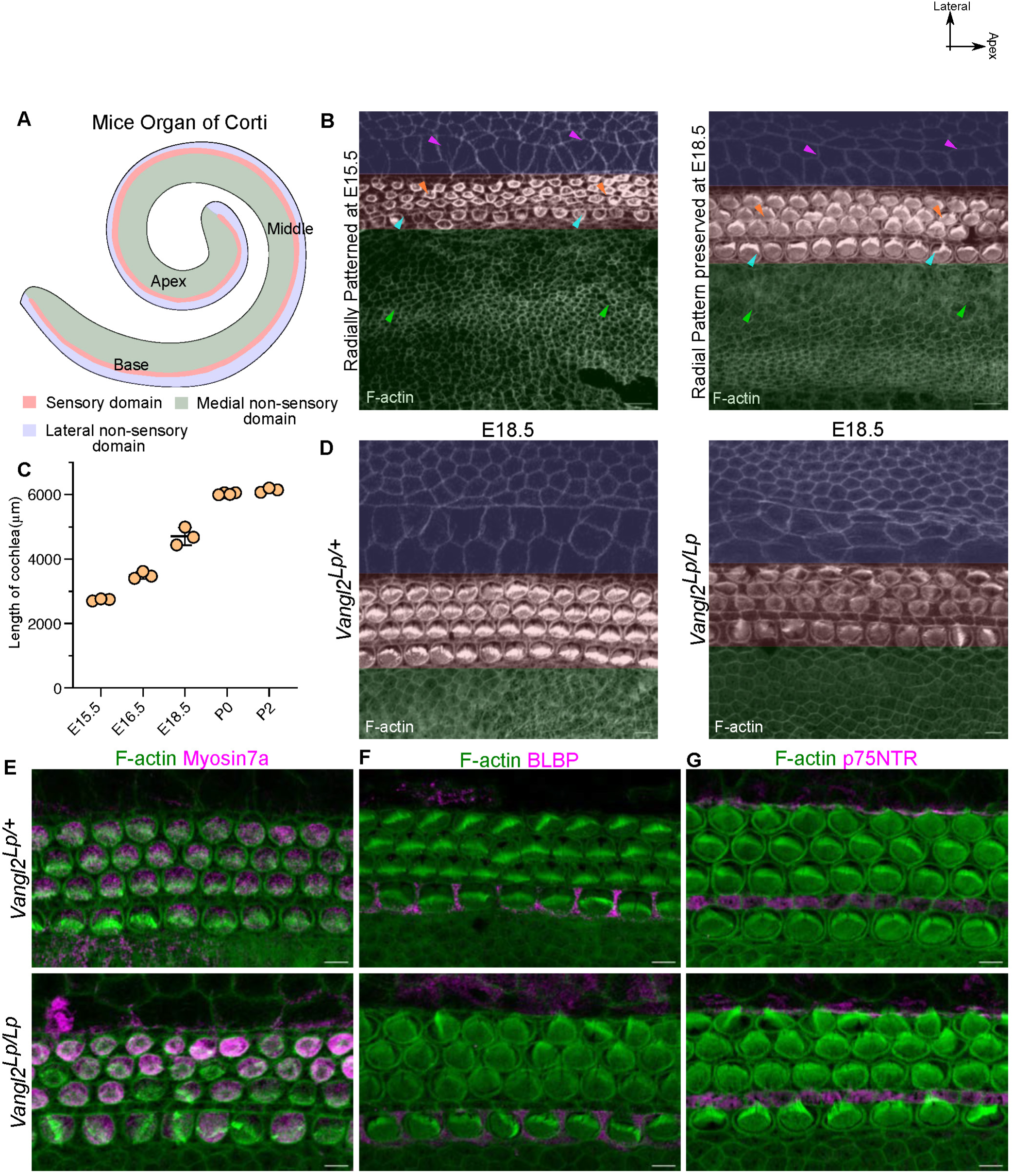
Cochlea undergoes pattern preserving convergent extension movement during development. A. Schematic of spiral-shaped mouse organ of Corti (OC) representing KÖlliker’s organ (medial non-sensory domain), sensory domain and lateral non-sensory domain. B. Base part of OC from E15.5 and E18.5 stained for F-actin. Green overlay and arrowhead indicate KO domain, red overlay indicates sensory domain and blue overlay indicates lateral non-sensory domain. Blue arrowhead indicates medial and orange arrowheads indicate lateral sensory domain, magenta arrowheads indicate Hensen’s cell. C. Length of cochlea from E15.5 to post-natal day 2. N=3 embryos for each stage. D. Base of E18.5 OC from heterozygous (*Vangl2 ^Lp/+^*) and homozygous (*Vangl2 ^Lp/Lp^*) looptail mutant stained for F-actin (grey). Overlay indicates domains as B. N=8 E. Base of E18.5 OC from heterozygous (*Vangl2 ^Lp/+^*) and homozygous (*Vangl2 ^Lp/Lp^*) looptail mutant stained for F-actin (green) and Myosin 7a (magenta and grey). N= 4. F. Base of E18.5 OC from heterozygous (*Vangl2 ^Lp/+^*) and homozygous (*Vangl2 ^Lp/Lp^*) looptail mutant stained for F-actin (green) and BLBP (magenta and grey). N= 4. G. Base of E18.5 OC from heterozygous (*Vangl2 ^Lp/+^*) and homozygous (*Vangl2 ^Lp/Lp^*) looptail mutant stained for F-actin (green) and p75NTR (magenta and grey). N= 4. Scale Bar: 5µm and 10µm in B and D. Image orientation: Top is lateral, Right is Apex.

From 15.5 to E18.5, the cochlea elongates from 2734±33µm to 4702±27µm (Fig. 1C, S1E). Previous studies have shown cell growth, migration, intercalation and tissue scale convergent-extension movements drive this elongation [13–17, 20]. Such movement should be expected to disrupt the organisation established at E15.5 [21]. However, they do not. Immunostaining the E18.5 OC for Myosin 7a, BLBP, p75NTR, revealed domain organisation was maintained during CE-mediated cochlear elongation (Fig. 1B, S1B-D).

While previous studies have investigated the organisation of the medial and lateral sensory domains when CE movements are perturbed, non-sensory domain organisation is unclear. Mice mutant for the core PCP protein, Vangl2, show defects in HC planar cell polarity and convergence and extension (CE) movements [22–25]. We thus assessed the organisation of non-sensory domains in these mutants. Homozygous *looptail* mutants of Vangl2 (referred as *Vangl2^Lp/Lp^*) have a cochlea 2/3^rd^ the length of littermate controls (*Vangl2^Lp/+^*5010±41µm and *Vangl2^Lp/Lp^* 3238±185µm) (Fig. S2A,B). Immunostaining for molecular markers for HCs, IPhC, inner pillar cells and the distinction in the morphological features of non-sensory cell types (Fig. 1D-G) revealed the relative position of cell types and domain organisation is maintained in the *Vangl2^Lp/Lp^* mutants with defects in convergent extension (Fig. S2C-E). This suggests a mechanism to maintain the integrity of domains during cochlear elongation.

### Adhesion Code defines compartments in the OC

Studies of mixed cultures of adhesion-molecule expressing cells and on gastrulating amphibian embryos and the fish neural tube have shown that the differential expression of adhesion molecule could segregate cells into domains [26–29]. The differential expression of cadherins and nectins has been described in sensory domain of OC, [9, 14, 19, 30, 31] leading us to characterise their expression across the entire OC. We focused on the expression of 3 cadherin family members, Cdh1, Cdh2 and Cdh4 (E-, N- and R-cadherin respectively), as well as nectin-1 and −2, and the tight junction protein ZO-1.

At E18.5, ZO1 and Nectin 2 are localised on the apical junctions of all the cells of the OC (Fig. S3A,B). Nectin 1 is expressed only on the HC-SC junctions (Fig. S3C). The expression of Cdh1 was found in the junctions of cells in the lateral domain of the OC encompassing lateral sensory and non-sensory domain. A faint expression of Cdh1 could also be seen in a subset of Kölliker’s organ cells bordering the medial sensory domain (Fig. S3D).

Cdh2 was expressed in the medial sensory and non-sensory domains (Fig. S3E). The expression of Cdh4 was similar to Cdh2, except expression was not detected in IPhC (Fig. S1F). This super-imposition of combinatorial expression of adhesion molecules and domains in OC suggested an adhesion-based segregation of cells in OC (Fig. S3G).

To ask when the putative adhesion code was established, we stained E14.5 cochlea for Cdh1, 2 and 4 before the onset of HC differentiation [32]. We find that even at E14.5, the expression of Cdh1, 2 and 4 was spatially restricted. Cdh1 shows expression in the lateral regions of the OC, with a faint expression detected on the medial edge of the putative KO domain (Fig. 2A). Cdh2 and Cdh4 are restricted to the medial OC (Fig. 2B, C). The presence of combinatorial cadherin expression before the onset of differentiation and its persistence during convergent extension-mediated elongation suggested its role in maintaining domain integrity during cochlear morphogenesis (Fig. 2D-E).

**Figure 2.**
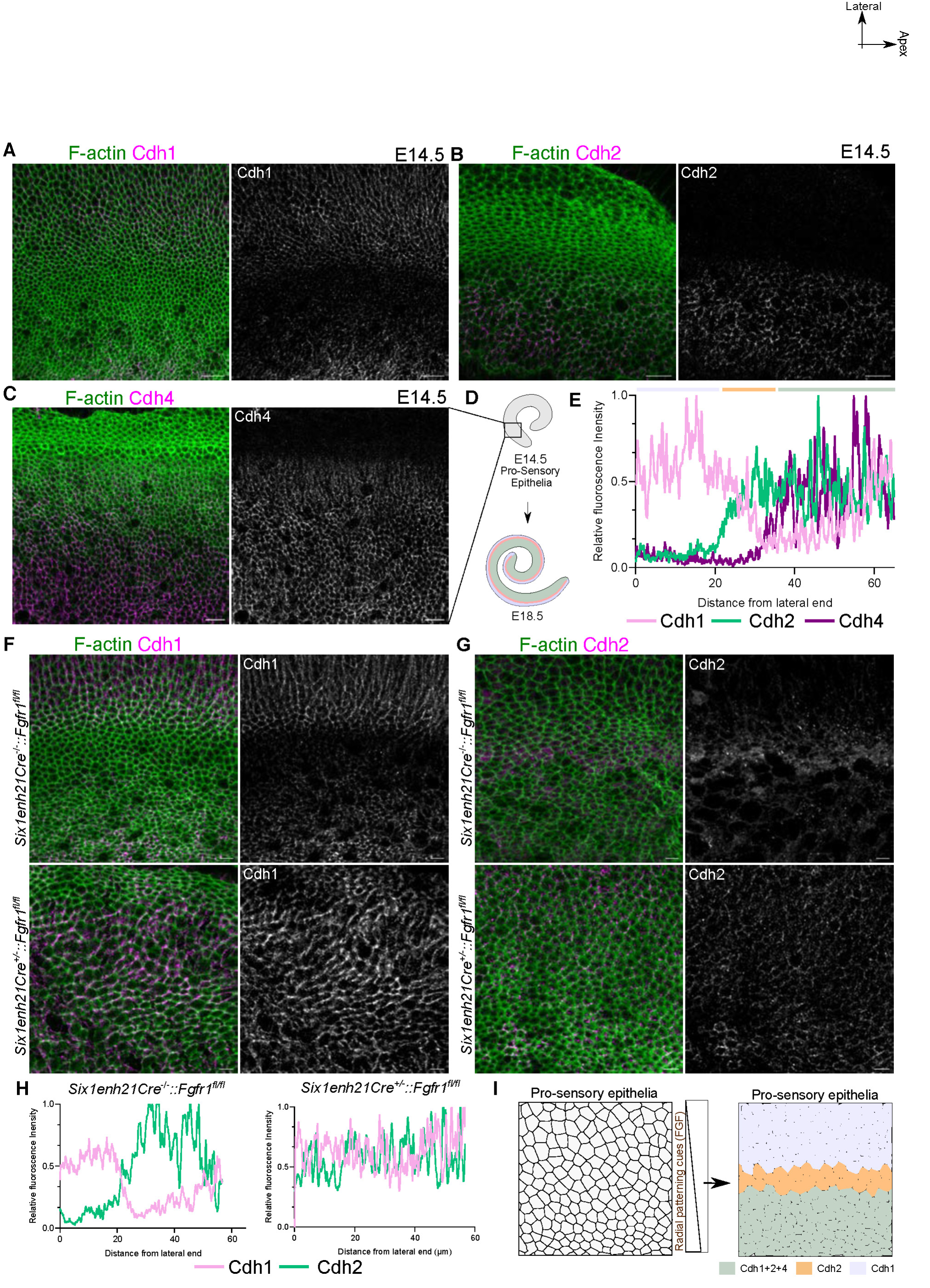
Radial patterning cues regulate development of adhesion-code based compartments. A. E14.5 OC stained for F-actin (green) and Cdh1 (magenta and grey). B. E14.5 OC stained for F-actin (green) and Cdh2 (magenta and grey). C. E14.5 OC stained for F-actin (green) and Cdh4 (magenta and grey). Image is medially shifted to show the expression in the KO. D. Schematic representing the pro-sensory epithelia at E14.5 develops into E18.5 OC. E. Relative fluorescence intensity of Cdh1 (pink), Cdh2 (green) and Cdh4 (purple) along the medio-lateral axis of OC at E14.5, representing lateral domain with only Cdh1 (lilac), a Cdh2 expressing domain (yellow) and a medial domain expressing all three cadherins (light green). F. E14.5 OC from control (*Six1enh21Cre^−/−^::Fgfr1^fl/fl^*) and fgfr1 mutant (*Six1enh21Cre^+/−^ :: Fgfr1^fl/fl^)* cochlea stained for F-actin (green) and Cdh1 (magenta and grey). N=3 embryos. G. E14.5 OC from control (*Six1enh21Cre^−/−^::Fgfr1^fl/fl^*) and fgfr1 mutant (*Six1enh21Cre^+/−^ :: Fgfr1^fl/fl^)* cochlea stained for F-actin (green) and Cdh2 (magenta and grey). N=3 embryos. H. Relative fluorescence intensity of Cdh1 (pink), Cdh2 (green) along the medio-lateral axis of OC at E14.5 from control (*Six1enh21Cre^−/−^::Fgfr1^fl/fl^*) and mutant (*Six1enh21Cre^+/−^:: Fgfr1^fl/fl^)* cochlea. I. Schematic representing the reciprocal expression of cadherins driven by radial patterning cues. Scale Bar: 10µm Image orientation: Top is lateral, Right is Apex.

To test this, we sought to disrupt the differential expression of cadherins. Work on zebrafish neural tube implicated a role for the morphogens involved in cell fate patterning in regulating adhesion molecule expression [29]. In mouse OC, FGF signalling, through the Fgfr1, has been shown to influence radial patterning (Huh et al., 2011; Ono et al., 2014) we drove the deletion of Fgfr1 from the inner ear using an inner ear-specific Cre (Six1enh21Cre). While E14.5 controls (*Six1enh21Cre^−/−^::Fgfr1^fl/fl^*) showed the combinatorial expression of Cdh1 and Cdh2, in mutant (*Six1enh21Cre^+/−^::Fgfr1^fl/fl^*^)^ cochlea, the patterned expression of Cdh1 and Cdh2 OC was absent (Fig. 2F-I). Immunostaining for Cdh1 and Cdh2 at E18.5 revealed that this misexpression is still evident and correlated with the sporadic presence of HCs in the medial and lateral non-sensory domains (Fig. S4), suggesting a reduction in the domain integrity.

To ask if the control of Cdh1 and Cdh2 expression by FGF signalling was independent of the cell fate specification, we used explant cultures of E16.5 cochlea (Fig. S5A). At this stage cell fates are already established, and CE-mediated elongation is still in progress. E16.5 cochlea explants treated with SU5402 for 12 hours, showed normal expression of p75NTR and BLBP suggesting negligible impact on cell fate (Fig S5B, C). However, staining these FGF inhibited cochlea for Cdh1 and Cdh2 showed misexpression of Cdh2 (Fig. S5D-F). Further, FGFR inhibited cochlea, showed ingression of pillar cells, frequent HC-HC contacts and drifting of HCs into medial and lateral non-sensory, suggesting perturbation of domain organisation (Fig S6A-C).

We next asked if the cadherin code itself was responsible for the preservation of domain organisation. Using function blocking antibodies, we asked if by perturbing cadherin interactions, we could perturb OC domains. To first establish that these antibodies were able to recognize Cdh1 and 2 in the cochlea, we first cultured E16.5 cochlea explants in presence of 7D6, an Cdh1 blocking antibody, or 6B3 which blocks Cdh2 interactions, for one hour and then stained with the respective secondary antibodies [33, 34]. We observed 7D6 and 6B3 antibodies localized to the lateral and medial regions of OC, similar to the expression of Cdh1 and Cdh2 (Fig S3D, E and Fig. S6D, E). We next cultured E16.5 cochlea explants in presence of either media containing BSA, 6B3 or 7D6 for 12 hours. Explants treated with the Cdh2 blocking antibody, 6B3 showed perturbations of the medial sensory and non-sensory domains, with IHCs drifting into the medial non-sensory domain and frequent HC contacts (Fig. S6F-H). Cochlea explants cultured in the presence of the Cdh1-blocking antibody, 7D6 led to the disruption of organisation in the lateral sensory domain, with negligible impact on medial non-sensory and sensory domain (Fig. S6F-H).

The similarity in SU5402 and 7D6 treated OC and the sustained mis-expression of cadherins in FGF-inhibited cochlea suggests FGF-driven combinatorial expression of cadherins underlines the domain integrity during convergent extension. The maintained expression of Cdh1,2 and 4 in Vangl2^Lp/Lp^ mutants with defective convergent extension further supported this inference (Fig. S7).

### Adhesion-code ensures discrete organisation of compartments during convergent extension

Previous work on fly imaginal discs, and the hindbrain, trachea, ovarian follicles, and somites [35–37] have provided evidence that cells within a compartment undergo distinct cell behaviours. These allow discrete tissue morphogenesis. To ask if adhesion-code based compartments in the cochlea also show distinct patterns of organisation, we analysed their development at two scales. The first is to ask how compartment shape changes, which we refer to as domain organisation. The second is to understand how the organisation of cells within each domain changes, referred to as cellular organisation. Previous studies have established that the sensory domain elongates (Fig. S8A-C) along the proximal-distal axis of OC while HCs and SCs within the domain migrate and intercalate to form a hexagonal lattice-like organisation between E15.5 and E18.5 [13, 38]. Hence, we focused on the non-sensory domains.

Kölliker’s organ is found medial to the sensory domain. Consistent with the increase in cochlear length, the KO also elongates twice its length along the base-apex axis of the cochlea (2734±33µm to 4702±27µm).). The extension of KO length is accompanied by a decrease in its width at the base and middle turn of the cochlea between E15.5 and E18.5 indicative of convergent extension (Fig. 3A, B). To understand the changes in cellular organisation, we investigated changes in apical cell shape and size. At E15.5, most cells of the KO have varied apical surface area and a roughly symmetrical aspect ratio (AR=0.87±0.12) closer to 1 (Fig. 3A, C). However, a small population of cells show an asymmetric elongation with their long axis aligned perpendicular to the mediolateral axis of the OC (Fig. 3D). By E18.5, two populations of cells are visible in the KO. Those at the medial edge of the KO (mKO), have a smaller apical surface area and are elongated, with the long axis parallel to the tissue axis (Fig. 3A, C, D). At the lateral KO (lKO), closer to the sensory domain, cells have a larger surface area and are elongated orthogonal to the tissue axis (Fig. 3A, C, D). On the lateral non-sensory domain, the apical surface area and the shape index (perimeter/sqrt of area) for both Hensen’s and Claudius cells increased between E15.5 and E18.5 (Fig. 3E, F and Fig.S8D, E).

**Figure 3.**
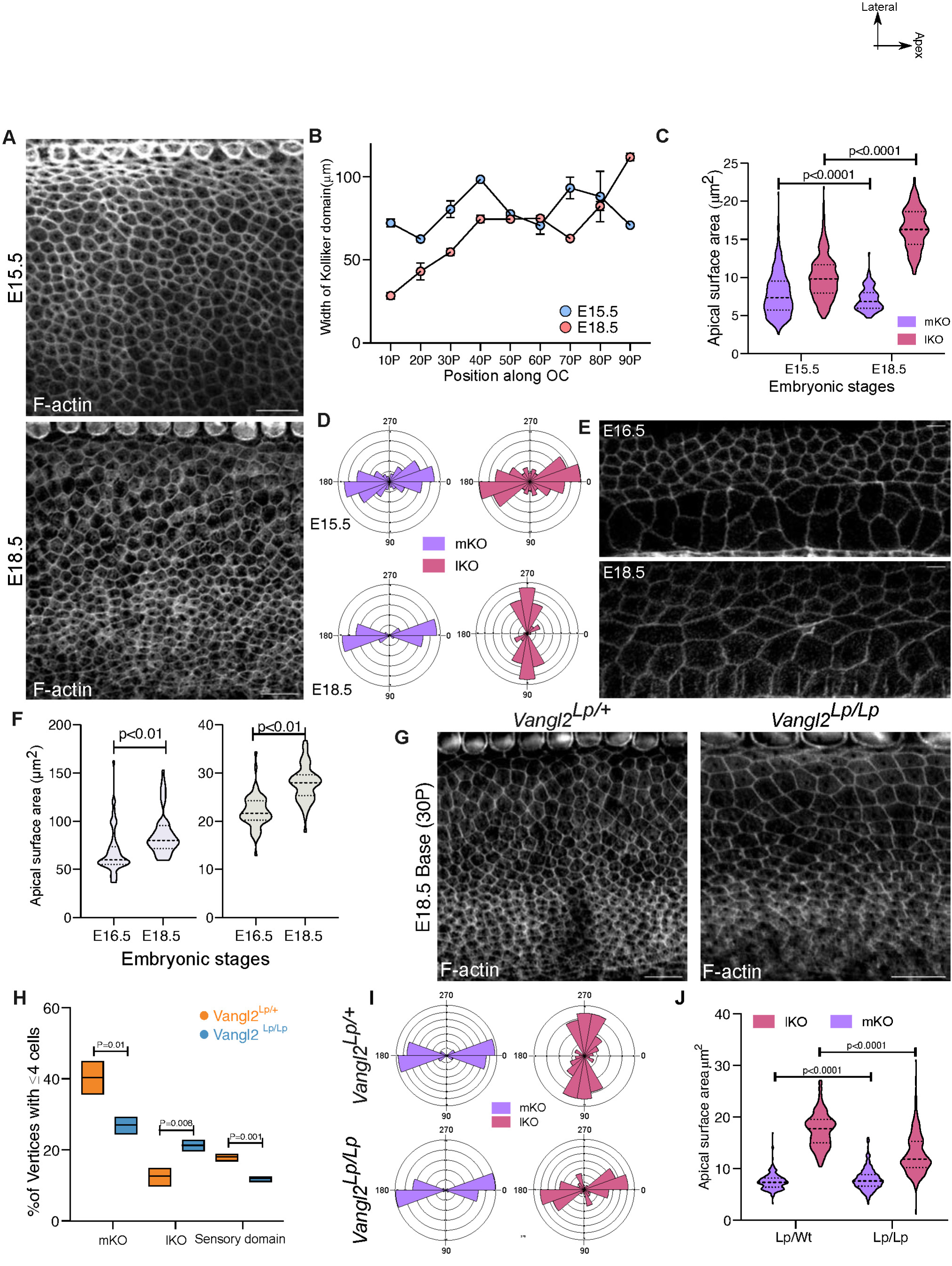
Planar cell polarity cues regulate organisation of each domain. A. Base region (30P) of OC from E15.5 and E18.5 stained for F-actin (grey) representing the KO domain. B. Width of the Kölliker’s organ along the OC at E15.5 and E18.5. N=4 cochlea for each stage. C. Apical surface area of mKO and lKO at E15.5 and E18.5 at basal region (30P). N= 378/303 (mKO/lKO, E15.5) and 283/298 (mKO/lKO, E18.5); 4 cochlea each stage. D. Rose stack plot represents the elongation cell axis at mKO and lKO at E15.5 and E18.5. N= 378/303 (mKO/lKO, E15.5) and 283/298 (mKO/lKO, E18.5); 4 cochlea each stage. E. Base region (30P) of OC from E15.5 and E18.5 stained for F-actin (grey) representing the lateral non-sensory domain containing Hensen’s cell and Claudius cells. F. Apical surface area of Hensen’s and Claudius cells at base region of OC at E16.5 and E18.5. N=64/60 for Hensen and 57/59 for Claudius (E16.5/E18.5). G. Base region (30P) of E18.5 OC from heterozygous (*Vangl2 ^Lp/+^*) and homozygous (*Vangl2 ^Lp/Lp^*) looptail mutant stained for F-actin (grey). H. Percentage of vertices with 4 or more cells for mKO, lKO and sensory domain in OC from heterozygous (*Vangl2 ^Lp/+^*) and homozygous (*Vangl2 ^Lp/Lp^*) looptail mutant at E18.5. N= 3 cochlea each: 250/622 mKO; 113/923 lKO; 155/880 sensory in Het and 249/921 mKO; 261/1215 lKO; 131/1079 sensory in Homo. (4 or more cell vertices /Total vertices) I. Rose stack plot representing the axis of cell elongation for mKO and lKO cells at the base region (30P) of E18.5 OC from heterozygous (*Vangl2 ^Lp/+^*) and homozygous (*Vangl2 ^Lp/Lp^*) looptail mutant. N=218/224 (mKO/lKO, Het) and 234/198 (mKO/lKO, Homo). J. Apical surface area of mKO and lKO cells at Base region (30P) of E18.5 OC from heterozygous (*Vangl2 ^Lp/+^*) and homozygous (*Vangl2 ^Lp/Lp^*) looptail mutant. N=218/224 (mKO/lKO, Het) and 234/198 (mKO/lKO, Homo). Scale Bar: 10µm Unpaired T-test, Image orientation: Top is lateral, Right is Apex.

Our data suggest that the non-sensory domains of OC, similar to the sensory domains, undergo distinct and discrete re-organisation during cochlear morphogenesis, a hallmark of compartment behavior. To ask if disrupting compartments also leads to a disruption in both domain and cellular organisation in the non-sensory compartments, we used mutants in which the adhesion code, and thus compartment cohesion, was disrupted. In Fgfr1 mutants, we observed a reduction in the differences between the apical surface area between the cells of the mKO and lKO (Fig. S8F, G). The cells from mutants also showed a reduction in circularity (Fig. S8F, G). Further the Hensen’s cells from mutants showed an increase in the shape index with similar apical surface area (Fig. S8I-K), suggesting a decrease in the compartment specific organisation.

### PCP regulated NMII-activity drives distinct cellular organisation

To understand the mechanisms behind discrete cellular organisation in compartments of the OC, we first looked at proliferation. We injected EdU into pregnant females at E13.5, E15.5, E16.5 and E18.5 and fixed embryos 6 hours after injection (Fig. S9A). At E13.5 (+6 hours), we observed the entire OC incorporated EdU (Fig. S9B), similar to previous observations [17]. By E15.5, the number of EdU positive cells decreased to less than 10% at the base. By E16.5, proliferation had ceased and remained so till E18.5 (Fig. S9B). As the KO increases in length between E15.5 and E18.5, we conclude that there is a limited contribution by proliferation. We next asked if cellular rearrangements could contribute to compartment reorganisation. In epithelia, cellular rearrangements result from neighbour exchange with an obligatory intermediate step where 4 or more cells meet at a vertex. We thus assessed the number of 4-cell vertices between E15.5 and E18.5. At E15.5, 33% of all vertices in the sensory compartment have 4 or more cells. This decreases significantly such that by E18.5 only 20% of vertices are made up of 4 or more cells (Fig. S9D).

Similarly, in KO domain, mKO showed 40% of vertices to have 4 or more cells suggesting a higher rate of cellular reorganisations. In the lKO, only 20% of vertices show 4 or more cells, suggesting a lower rate of reorganisation compared to the mKO. While the proportion of vertices with 4 or more cells decreased in the lKO by E18.5, the proportion of vertices with 4 or more cells in the mKO at E18.5 remained equivalent to the numbers observed at E15.5 (Fig. S9E). This data suggests cells in KO domain undergo cell rearrangement, higher at medial edge compared to the lateral edge. In the absence of cell division, we hypothesized this cellular rearrangement to drive cochlear morphogenesis.

To test this, we sought to perturb the process of cellular re-organisation by disrupting activity of acto-myosin complex, essential for cellular intercalations. Non-muscle myosin forms the motor component of the acto-myosin complex. The motor activity of NMII is regulated by the phosphorylation status of its regulatory light chain (RLC). Thus, we first immuno-stained OC for mono and di-phosphorylated forms of RLC. At E18.5 p-RLC is expressed on all junctions (Fig. S1A). While the pp-RLC localised at the medial edge of OHC-DC junctions (Fig. S10B, B’), IPhC-Pillar cells junctions (Fig. S14C, D), and in the KO compartment, it was found along the junctions of the long-axis of KO cells (Figure S14B). To test their role in morphogenesis of cochlea, we used our ex-vivo explant method to culture E16.5 cochlea for 8 hours, in the presence or absence of ML7 (which inhibits Myosin Light Chain Kinase (MLCK)) [39, 40]. In MLCK-inhibited OC, we observed a decrease in apical surface area and circularity of OHC compared to the control samples (Fig. S10C-H), suggesting a decrease in spatial organisation within the sensory domain. Previous work on avian auditory epithelia have shown the spatial organisation of HC is coupled to alignment of HC polarity to the tissue axis [12]. Similarly, we observed a decrease in the alignment of HC polarity for IHC and OHC in the MLCK inhibited cochlea (Fig. S10E, F). In addition, the differences in the apical surface area and the elongation axis of mKO and lKO cells was also reduced in the MLCK-inhibited OC (Fig. S10I-K). This data suggested NMII driven neighbor-exchange drives reorganisation of sensory and non-sensory domains during development.

To further understand this organisation, we decided to understand how NMII activity is regulated in each compartment. Previous studies including our work on avian auditory epithelia have shown planar cell polarity cues through Vangl2 regulates RLC phosphorylation [12, 41–43]. In mouse the expression of Vangl2 largely overlapped with the expression of pp-RLC (Fig. S11A, B). Hence, we used Vangl2 ^Lp/Lp^ mutant which, as previously reported, shows a reduction in alignment of HC polarity (Fig. S11C, D) [15, 22]. pp-RLC shows a down-regulation on the junctions of both sensory and non-sensory compartments in Vangl2 ^Lp/Lp^ mutants, while the p-RLC was comparable to the littermate control (Fig. S11E-H). At the scale of the domain, the Vangl2 ^Lp/Lp^ mutants showed a decrease in the width of both KO and sensory domain at the base and middle turn of OC (Fig. 3G and Fig. S11I-J). Further, Vangl2 ^Lp/Lp^ mutants at E18.5, also showed a significant decrease in the number of 4 cell vertices in both mKO and sensory domain, suggesting a decrease in cell intercalation (Fig. 3H). Interestingly, mutants showed a significant increase of 4 cell vertices in the lKO. The differences in the apical surface area, preferential axis of elongation for mKO and lKO cells were also reduced in the mutants (Fig. 3G, I, J). The regulation of RLC phosphorylation by Vangl2 and the similarity of Vangl2 ^Lp/Lp^ mutants with MLCK inhibited OC, suggests Vangl2 regulated NMII activity may govern the organisation of not only the sensory compartments of the OC but also the non-sensory compartments.

### Compartment-intrinsic remodeling has extrinsic effects on cellular organisation

To understand the effect of disrupting cell and domain organise in a compartment, we used the Cre-LoxP system to delete Vangl2 from individual domains. We generated mice carrying one copy of the looptail mutation of Vangl2 and the other allele, the conditional Vangl2 mutant, where the coding sequence is flanked by loxP sites. We referred to this mouse as *Vangl2^lp/fl^*.

Previous studies using an inducible CreER line driven by Neurogenin, identified expression in cells within the KO, as well as in the cochleovestibular ganglion. Importantly, cells within the sensory compartment of the OC were not labelled [44]. We first verified that a non-inducible Cre-line, Ngn1^457^-Cre could recapitulate this expression domain [45]. Ngn1^457^-Cre males were crossed with Ai14 (ROSA26-TdTomato) females to generate Ngn1^457^-Cre::Ai14. OC from E18.5 embryos were dissected and imaged. Similar to Ngn1-CreER, we found that Ngn1^457^-Cre could drive recombination in around 10% of cells located throughout the KO (Fig. 4A, B). Recombination was not detected in the sensory compartment. At E18.5, the cochlea from the experimental animal (*Ngn1^457^-Cre^+/−^:: Vangl2^lp/fl^*) showed a reduction in difference of apical surface area between mKO and lKO cells compared to the littermate control ((*Ngn1^457^-Cre^−/−^:: Vangl2^lp/fl^*)) cochlea (Fig. 4C,D). Similarly, the long axis of lKO cells in (*Ngn1^457^-Cre^+/−^:: Vangl2^lp/fl^*) mutants was parallel to the OC tissue axis, compared to the orthogonal direction in the control cochlea (Fig. 4C,D). This suggests that a compartment-specific deletion of Vangl2 can perturb local cellular organisation.

**Figure 4.**
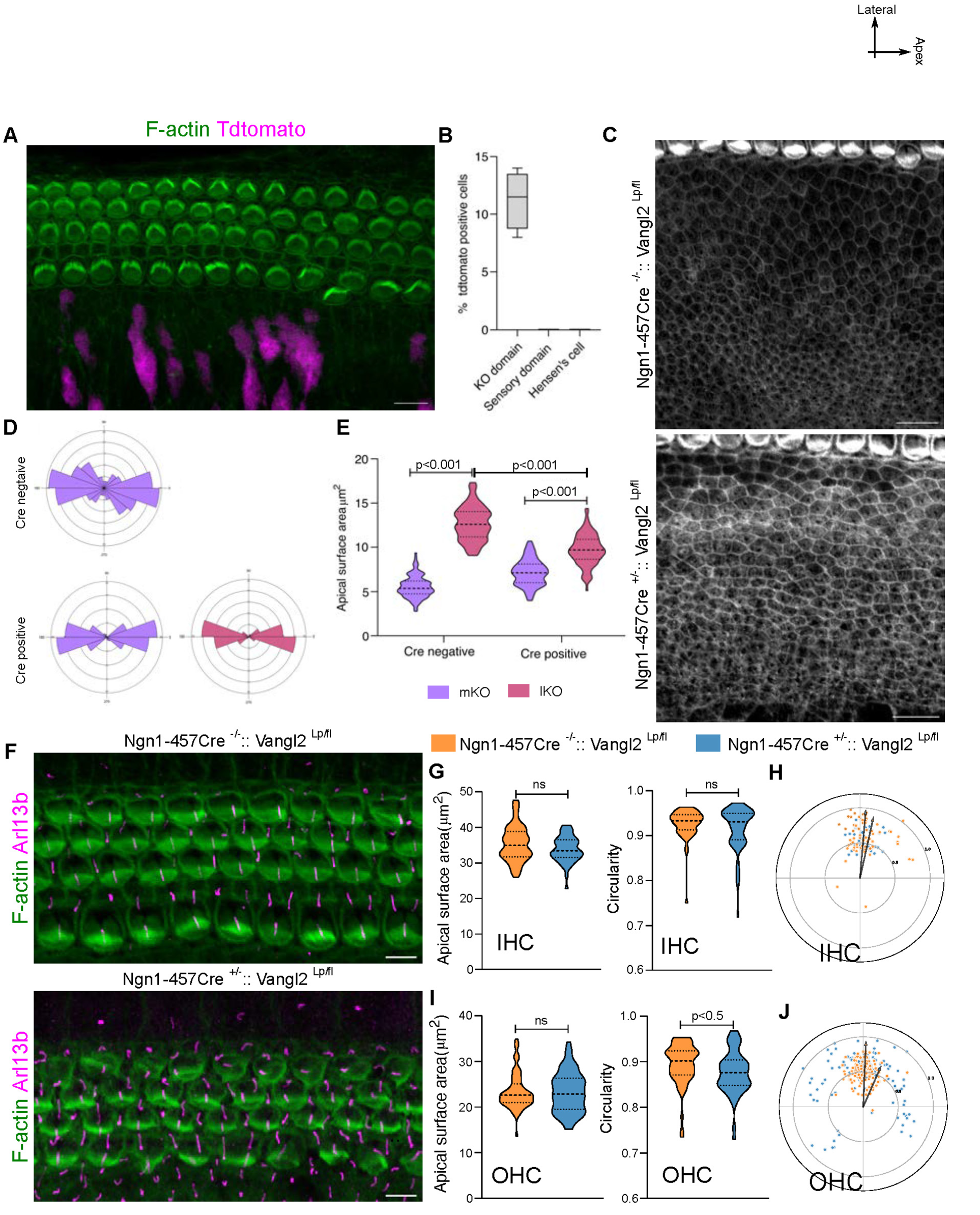
Deletion of Vangl2 from Kölliker’s organ disrupts organisation in lateral non-sensory domain. A. E18.5 OC from *Ngn1^457^-Cre^+/−^::Ai14*, stained with F-actin (green) showing cre mediated expression of Tdtomato (magenta) in Kölliker’s organ and not in sensory domain. B. Percentage of tdtomato positive cells from the *Ngn1^457^-Cre^+/−^::Ai14* cochlea in KO, sensory and lateral non-sensory domain. N=3 C. Kölliker’s organ from base region (30P) of OC from Cre negative control (*Ngn1^457^-Cre^−/−^:: Vangl2^lp/fl^*) and Cre positive mutants (*Ngn1^457^-Cre^+/−^:: Vangl2^lp/fl^*) stained for F-actin (grey). N=4 embryos. D. Rose stack plots representing the axis of cell elongation in medial and lateral KO cells from Cre negative control (*Ngn1^457^-Cre^−/−^:: Vangl2^lp/fl^*) and Cre positive mutant (*Ngn1^457^-Cre^+/−^:: Vangl2^lp/fl^*) OC. N= 190/202 for Control and 198/208 for Mutant (mKO/lKO). E. Apical surface area of medial and lateral KO cells from Cre negative control (*Ngn1^457^-Cre^−/−^:: Vangl2^lp/fl^*) and Cre positive mutant (*Ngn1^457^-Cre^+/−^:: Vangl2^lp/fl^*) OC. N= 190/202 for Control and 198/208 for Mutant (mKO/lKO). F. Base region (30P) of OC from Cre negative control (*Ngn1^457^-Cre^−/−^:: Vangl2^lp/fl^*) and Cre positive mutant (*Ngn1^457^-Cre^+/−^:: Vangl2^lp/fl^*) stained for F-actin (green) and Arl13b (magenta). G. Apical surface area and Circularity of IHC of Cre negative control (*Ngn1^457^-Cre^−/−^ :: Vangl2^lp/fl^*) in orange and Cre positive mutant (*Ngn1^457^-Cre^+/−^:: Vangl2^lp/fl^*) in blue. N=92/96 (control/mutant). H. Polar coordinates representing the position of kinocilia from IHC of Cre negative control (*Ngn1^457^-Cre^−/−^:: Vangl2^lp/fl^*) in orange and Cre positive mutant (*Ngn1^457^-Cre^+/−^:: Vangl2^lp/fl^*) in blue. N= 66/66 (control/mutant). I. Apical surface area and Circularity of OHC of Cre negative control (*Ngn1^457^-Cre^−/−^ :: Vangl2^lp/fl^*) in orange and Cre positive mutant (*Ngn1^457^-Cre^+/−^:: Vangl2^lp/fl^*) in blue. N= 143/151 (control/mutant). J. Polar coordinates representing the position of kinocilia from OHC of Cre negative control (*Ngn1^457^-Cre^−/−^:: Vangl2^lp/fl^*) in orange and Cre positive mutant (*Ngn1^457^-Cre^+/−^:: Vangl2^lp/fl^*) in blue. N= 113/121 (control/mutant). Scale Bar: 10 µm in A, C and 5µm in F Unpaired T-test, ns=P>0.05, non-significant. Image orientation: Top is lateral, Right is Apex.

We next asked if local perturbations in organisation may result in non-local perturbations in other compartments. The sensory domain, comprises of the medial and lateral sensory compartments. We did not observe significant defects in polarity, apical surface area or circularity in the immediately adjacent medial sensory compartment (Fig. 4F-H). In contrast, while surface area was unaffected in the lateral sensory compartment, we noted a decreased alignment of planar polarity of OHC3 in *Ngn1^457^-Cre^+/−^:: Vangl2^lp/fl^*mutants (Fig. 4F, I, J) and a reduction in the circularity of OHCs. This suggested that disruption of *Vangl2* from the non-sensory KO compartment could influence the development of planar polarity in the lateral sensory compartment.

Previous work on the vestibular sensory epithelium [46], wing imaginal disc [47] and ommatidia [48], together with their mathematical description [49, 50], suggests that a local deletion of PCP cues from a group of cells can also affect the polarity of neighbouring cells, a phenomenon known as domineering non-cell autonomy. The effect decreases away from the site of deletion. However, in *Ngn1^457^-Cre^+/−^:: Vangl2^lp/fl^* mutants, only the polarity of OHC3 cells is affected (Fig. 4J). These are at a considerable distance from the KO. As polarity in IHC of the medial sensory compartment is more developmentally advanced than OHC [51], it is possible that any defects in IHC polarity could have been observed at an earlier developmental stage. However, even at E16.5, IHC showed no disruption in the alignment of planar polarity. In contrast, the 3rd row of OHC of the lateral sensory compartment still showed a small deviation in HC polarity alignment as compared to the littermate control (Fig. S12A). Our *Ngn1^457^-Cre* driven deletion of Vangl2 from KO domain also contains a *looptail* allele, which has been suggested to exert a dominant phenotype. Although the comparison with the *Ngn1^457^-Cre^−/−^:: Vangl2^lp/fl^* littermate controls suggest that the *looptail* allele does not show dominance, we assayed the phenotype of the *Vangl2^fl/fl^* mutant. Using the *Ngn1^457^-Cre^+/−^:: Vangl2^fl/fl^*, where both alleles are floxed out by Cre-mediated recombination we find, similar to *Ngn1^457^-Cre^+/−^:: Vangl2^lp/fl^,* that *Ngn1^457^-Cre^+/−^:: Vangl2^fl/fl^* also showed a perturbation of the polarity of 3^rd^ row of OHC (Fig. S12B). This suggests that the loss of OHC3 alignment results from the loss of cellular organisation in the KO compartment. This can neither be explained by domineering non-cell autonomy nor the dominant effect of *looptail* mutation. Instead, we suspected a long-range effect between compartments of the OC.

To ask whether the sensory domain also influences the development of the non-sensory domain, we used *Lgr5-CreERT2*, a tamoxifen-inducible Cre which, when induced at E15.5, drives recombination in the sensory domain but not in the KO (Fig. S12C, D). In the animals where recombination is induced at E15.5, we observed a reduction in planar polarity and circularity for HC at E18.5 (Fig. 5A-E). To ask if the KO is affected, we measured the apical surface area and the direction of the long axis in the lateral and medial KO. The mutant, tamoxifen-induced, KO compartment showed a slight decrease in the difference between the apical surface area of the mKO and lKO cells (Fig. 5F, G). Moreover, mutant cochlea showed no difference between the direction of the long axis of lKO cells (mediolateral to the tissue axis) and that of the mKO cells (parallel to the tissue axis) (Fig. 5F, H). This data suggests a reciprocal influence of the sensory domain on cellular organisation within the KO compartment (Fig. 5I).

**Figure 5.**
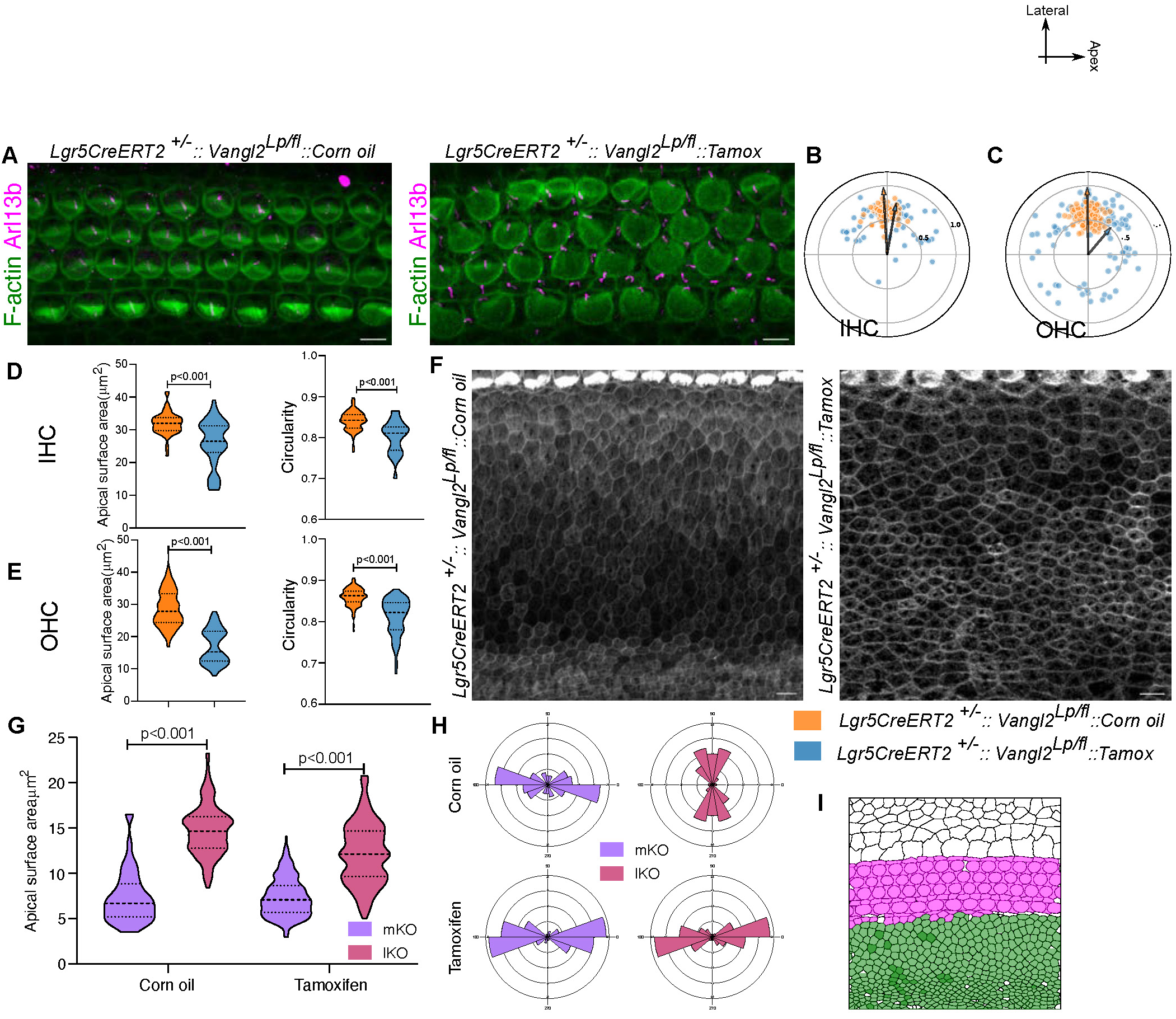
Deletion of Vangl2 from sensory compartments disrupts organisation in KO compartment. A. Base region (30P) of OC from corn oil injected control (*Lgr5CreERT2^+/−^:: Vangl2^lp/fl^::Corn oil*) and tamoxifen induced mutant (*Lgr5CreERT2^+/−^:: Vangl2^lp/fl^::Corn oil*) stained for F-actin (green) and Arl13b (magenta). B. Polar coordinates representing the position of kinocilia of IHC from corn oil injected control OC in orange and tamoxifen-induced mutant in blue. N= 66/66 (corn oil/Tamoxifen). C. Polar coordinates representing the position of kinocilia of OHC from corn oil injected control OC in orange and tamoxifen-induced mutant in blue. N=110/110 (Corn oil/Tamoxifen) D. Apical surface area and Circularity of IHC from corn oil injected control OC in orange and tamoxifen-induced mutant in blue. N= 89/99 (corn oil/Tamoxifen) E. Apical surface area and circularity of OHC from corn oil injected control OC in orange and tamoxifen-induced mutant in blue. N= 109/128 (corn oil/Tamoxifen) F. Kölliker’s organ from base region (30P) of OC from corn oil injected control OC and tamoxifen-induced mutant embryos stained for F-actin(grey). Scale Bar: 10µm G. Apical surface area of medial and lateral KO cells from corn oil injected control and tamoxifen-induced mutant OC. N=265/245 for Corn oil and 264/234 for Tamoxifen (mKO/lKO). H. Rose stack plots representing the axis of cell elongation in medial and lateral KO cells from corn oil injected control and tamoxifen induced mutant OC. N=265/245 for Corn oil and 264/234 for Tamoxifen (mKO/lKO). I. Schematic representing the expression of cre in lateral and medial sensory compartments (magenta) and hence the deletion of Vangl2, with the effect on the organisation in the KO (green). Scale Bar: 5 µm in A and 10 µm in F Unpaired T-test. Image orientation: Top is lateral, Right is Apex.

Our results show the deletion of Vangl2 from a compartment of OC, influences the organisation of other distant compartments. The influence of one compartment on the organisation of another suggested a mechanism that integrates and coordinates non-local cellular behaviour, which we referred to as compartment coupling.

### Mechanical origin of Coupling

There are potentially many mechanisms which could drive coupling among compartments. However, as we found this coupling in domain-specific mutants of Vangl2, and because Vangl2 regulates junctional mechanics, we hypothesized that coupling may have a mechanical component. To test this, we used a mutant of Vinculin, a protein important for transmission of forces between junctions [52]. We reasoned that by removing this protein, the mechanical coupling between compartments would be impaired (Fig. 6A). We used a conditional allele of Vinculin (*Vinc^fl/fl^*) and deleted this in the cochlea from E12.5 using Emx2-Cre. We could only obtain 3 experimental embryos out of 114 examined (see methods for detail). The OC from *Emx2-Cre^+/−^::Vinc^fl/fl^* showed disruption in the alignment of planar polarity in the lateral sensory compartment (Fig. 6B-F). In the KO, there is now a population of larger lKO cells abutting the medial sensory compartment (Fig. 6B, G, H) in mutants. The differences in the direction of the long axis of lKO and mKO cells are reduced in *Emx2-Cre^+/−^::Vinc^fl/fl^* (Fig. 6G, I). These data suggest the coupling between compartments is mediated, at least in part, through mechanics.

**Figure 6.**
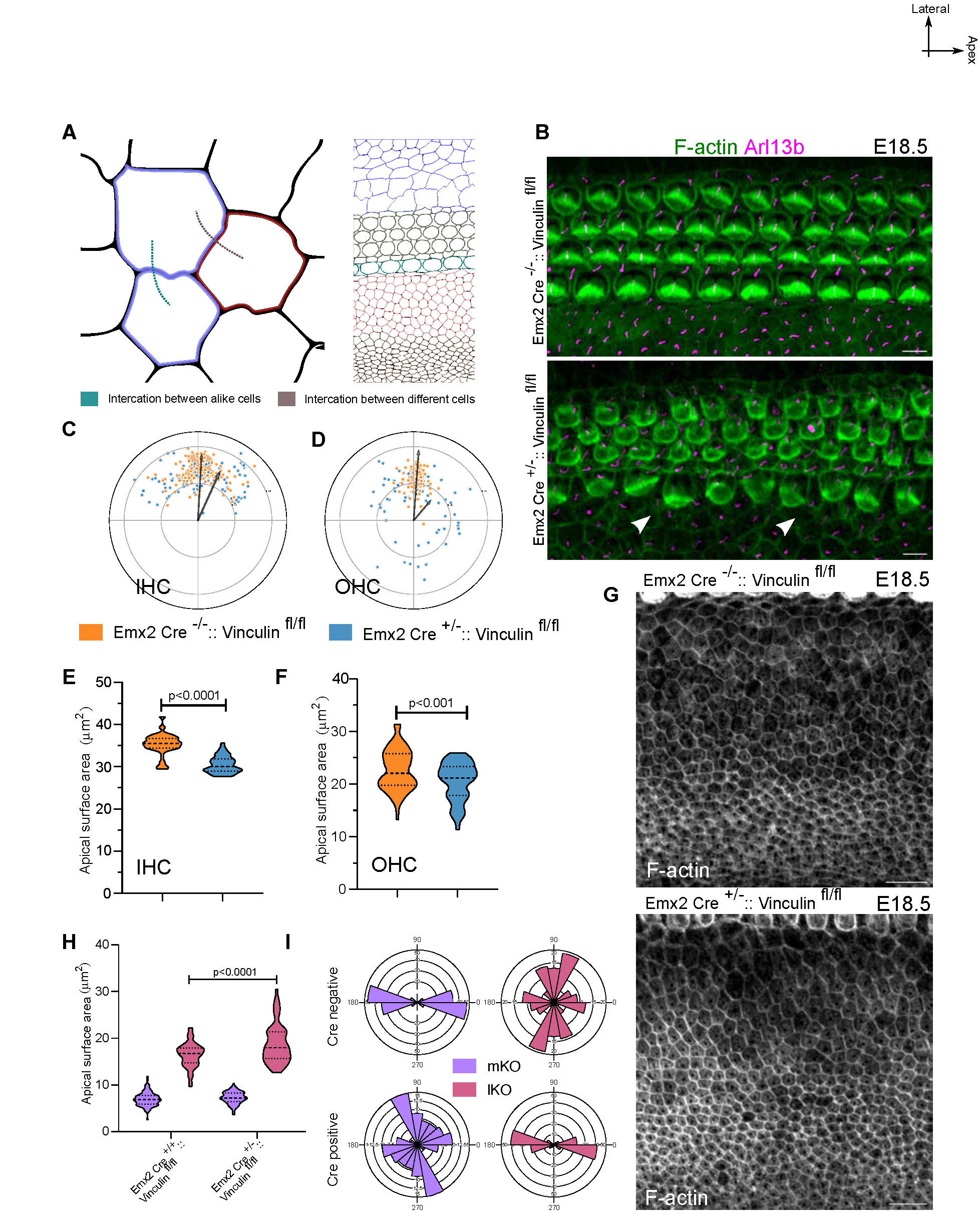
Disruption of Junctional force transmission complex protein, Vinculin, disrupts the organisation of OC. A. Schematic representing the communication between cells of OC with diverse mechanical properties. We consider the cells of OC are mechanically different and the communication can be between cells of same type and cells of different types. B. Base region (30P) of E18.5 OC from the control (*Emx2-Cre^−/−^:: Vinculin ^fl/fl^*) and mutant for vinculin (*Emx2-Cre^+/−^:: Vinculin ^fl/fl^*) stained for f-actin (green) and Arl13b (magenta). Arrow represents the disrupted organisation of Inner border cells. C. Polar coordinates representing the Kinocilium position of IHC from control (orange) and vinculin mutant (blue) at base region (30P) of OC. N=66/66 (control/Mutant) D. Polar coordinates representing the Kinocilium position of OHC from control (orange) and vinculin mutant (blue) at base region (30P) of OC. N= 68/68 (control/Mutant). E. Apical surface area of IHC from control (orange) and vinculin mutant (blue) at base region(30P) of OC. N=66/66 (Control/Mutant) F. Apical surface area of IHC from control (orange) and vinculin mutant (blue) at base region(30P) of OC. N=66/66 (Control/Mutant) G. KO of base region of OC (30P) from control and Vinculin mutant OC stained for F-actin (grey). H. Apical surface area of medial and lateral KO cells from base region (30P) of control and Vinculin mutant OC. N=145/165 (Control/Mutant) I. Rose stack plots representing the axis of cell elongation in medial and lateral KO cells from base region (30P) of control and Vinculin mutant OC. N=145/165 (Control/Mutant) Scale Bar: 5 µm in B and 10 µm in G Unpaired T-test. Image orientation: Top is lateral, Right is Apex.

## DISCUSSION

Using mouse auditory epithelia, we investigate a fundamental question in morphogenesis: How are developmental instructions coordinated across spatial scales so that tissues form shapes and patterns that are robust and reproducible? We demonstrate that the organ of Corti is segregated into compartments, adhesion-based spatially restricted developmental units, which allow for locally coordinated, discrete cell behaviours. Surprisingly, by using compartment-specific perturbations of cell mechanics, we find intra-compartment defects have effects on the cellular organisation of other compartments. We consider this as evidence of inter-compartment coupling and suggest that it provides a mechanism allowing the transmission of morphogenetic information across spatial hierarchies, ensuring fidelity in organisation.

Compartment coupling has been proposed as a mechanism to coordinate adjacent domains. This is the case during axial elongation [53], where force coupling ensures the coordination of neighbouring axial and paraxial mesoderm. Similarly, recent studies during hair follicle development have found coupling between adjacent compartments necessary for the invagination and polarity of the feather bud and hair follicle [54–56]. The behaviours between adjacent compartments can be considered linear, and studies from the coordination of the prechordal plate with the anterior neural plate in the zebrafish suggest that cells further from the compartment interface show a reduced coupling effect [57]. In contrast, the coupling shown in the mouse OC is both linear (between the adjacent sensory domain and the lKO) and non-linear, as the tissues effected are not adjacent. How does the non-linear misalignment of OHC3 emerge from perturbing KO? Previous studies have identified forces at the interface of the lateral non-sensory compartment (containing Hensens Cells) and the lateral sensory compartment (containing the OHCs) as a source of shear-induced crystallization that orders the HC-SC mosaic in the lateral sensory compartment [11]. It is possible that by disrupting KO organisation, this shear force is perturbed, with an effect on OHC ordering. Consistent with this is the effect of Vangl2 reduction from the KO reducing HC circularity in OHC.

Our data identifies compartments as mechanisms that permit distinct patterns of cellular organisation within an organ. Compartments bridge scales between embryonic fields and cells, and by coupling cellular behaviours between these compartments, they fine-tune positional information. Additionally, the coupling is likely contingent on the extent of intra-compartment organisation, adding a temporal aspect. Thus, it is probable that coupling not only organises developmental instructions spatially but also temporally. We suggest that compartments and their coupling provide a conceptual framework to spatially and temporally organise morphogenetic information relevant not only to the development of the inner ear but also, more generally, during organogenesis.

## METHODS

### Mice

#### Animal housing

All mice were housed at Animal Care and Resource Centre (ACRC) at NCBS, in accordance with Institutional Animal Ethics Committee guidelines.

#### Mouse strains

The supplementary table below provides information about the mouse strains used.

**Table S1:** Mouse strains used for experiments in this paper.

| Mouse strain | Identifier | Source |
| --- | --- | --- |
| Vangl2 (looptail) | JAX:000220 | Jackson Laboratory |
| Vangl2 Flox | JAX: 025174 | Jackson Laboratory |
| TdtomatoAi14 | JAX: 007914 | Jackson Laboratory |
| Fgfr1 Flox | NA | Ulla Pirvola 2002 |
| Emx2 Cre | NA | Jun Kimura 2005 |
| Sox10 Cre | JAX:025807 | Jackson Laboratory |
| Six1enh21 Cre | NA | Shigeru Sato 2012 |
| Ngn1<sup>457</sup> Cre | JAX:012859 | Jackson Laboratory |
| Lgr5-IRES-CREERT2 | JAX:008875 | Jackson Laboratory |
| Vinculin Flox | NA | Alice E. Zemljic-Harpf 2007 |

#### Genotyping

For maintaining stock animals of strains, an ear biopsy was collected from P21-P30 animals. Each biopsy was lysed in 150µl lysis solution (250ul of 1M NaOH, 2ul of 1M EDTA in 9.748 ml of autoclaved water) for 1 hour at 95°C in a 1.5ml microcentrifuge tube and then stored after adding 150µl neutralising solution (0.4ml of 1M Tris-Cl in 9.6ml of autoclaved water). 1µl of this solution was used as template for polymerase chain reaction (PCR) based genotyping. A 10µl of PCR mixture contained 5µl of Kappa-2X genotyping master mix (Cat# KK1024, Roche), 0.5µl each of reverse and forward primer and 3µl of nuclease free water. The PCR was performed as per manufacturer protocol and the oligonucleotide used as primers is listed in Table S2.

**Table S2:**
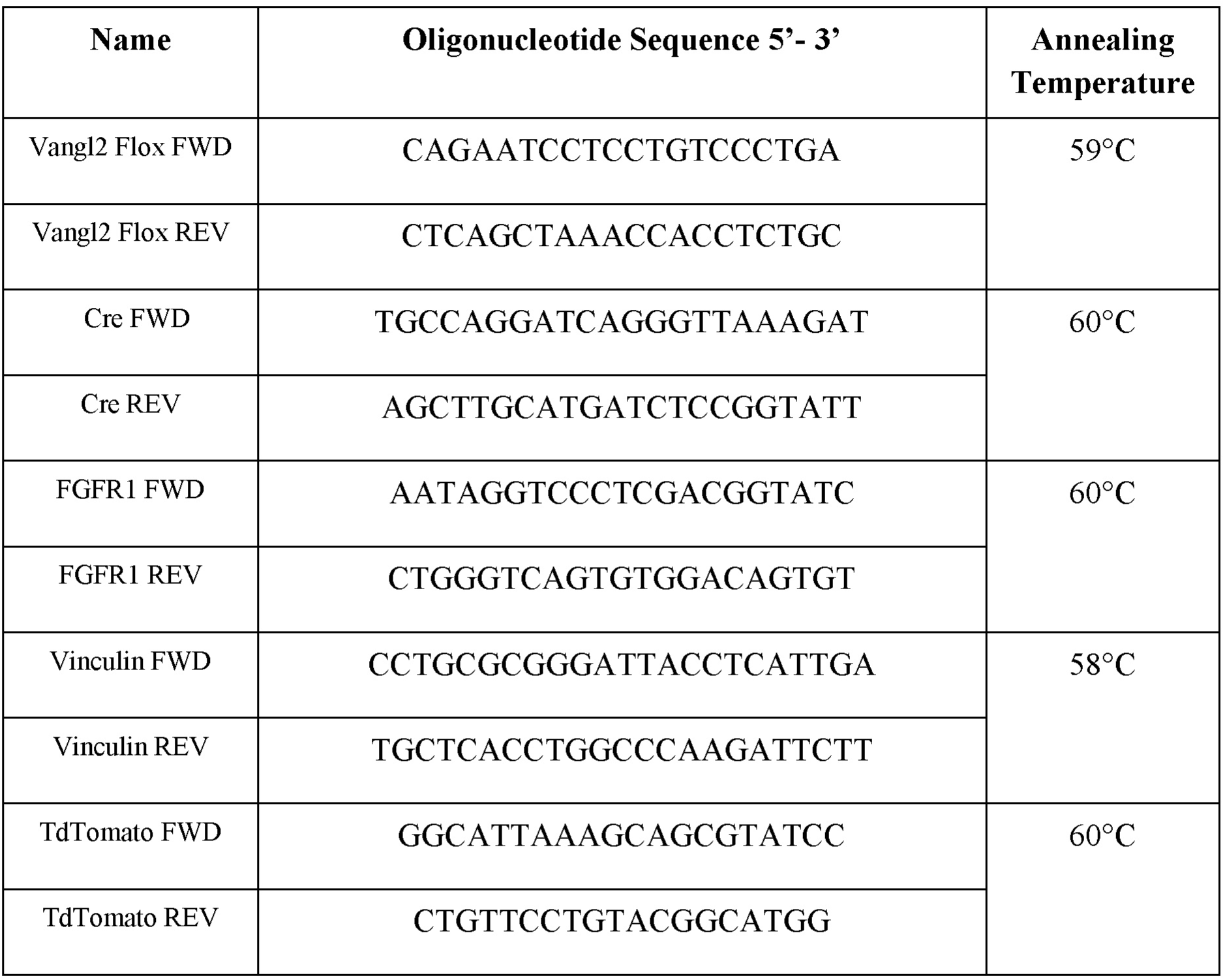
Oligonucleotide sequences used for PCR-based genotyping mouse strains.

*Looptail* mutants were identified with kinked to looped tails in heterozygous conditions and with an open neural tube in the homozygous conditions.

**Note:** The Vinculin flox mouse was crossed with both Sox10-Cre and Emx2-Cre to obtain Vinculin cKOs for analysis. However, we could not find any living embryos at E18.5 for *Sox10-Cre^+/−^::Vinc^fl/fl^*. Only 3 out of 114 *Emx2-Cre^+/−^::Vinc^fl/fl^* animals were obtained. The heterozygous *Emx2-Cre^+/−^::Vinc^+/fl^* developed a hydrocephalous-like phenotype.

#### Tamoxifen Injection

The age-appropriate female of the *Vangl2^fl/fl^* genotype was bred with *Lgr5-CreERT2 ^+/−^:: Vangl2^Lp/fl^* genotype males. Tamoxifen was dissolved in 90% corn oil and 10% ethanol to prepare a stock concentration of 10mg/ml. Pregnant females were weighed at the E15.5 stage using an electronic balance and were injected intraperitoneally with an appropriate volume of tamoxifen solution at 4.5mg/40g of body mass. As a control, corn oil and ethanol solution without tamoxifen is injected.

#### EdU injection

The age-appropriate female were injected with EdU (2.5mg/40g of animal weight) intraperitoneally using sterilized syringe and were dissected after 6 hours.

### Immunostaining

The inner ears from staged mouse embryos were dissected using a pair of forceps in ice-cold Phosphate Buffer Saline (PBS) and then fixed in 4% Paraformaldehyde (PFA) for 30 minutes to overnight according to the primary antibody, See Table S3. The inner ear is further dissected to expose sensory epithelia. The sensory epithelia are then permeabilized using 0.3% Tween-20 in PBS for 30 minutes at room temperature and blocked using blocking solution (5% heat-inactivated goat serum, 1% bovine serum albumin, 0.3% Tween-20 and PBS) for 1 hour at room temperature. Sensory epithelia are then incubated with primary antibodies diluted in a blocking solution (5% heat inactivated goat serum, 1% Bovine Serum Albumin, 0.3% Tween-20 in PBS) for overnight at 4°C. Sensory epithelia are then washed for 4 hours with a change of washing solution (0.3% Tween-20 in PBS) after every 30 minutes. The sensory epithelia are then incubated with secondary antibodies conjugated with Alexa fluor and phalloidin conjugated with Alexa fluor for 1 hour at room temperature and then washed using a washing solution for 3 hours with a change of washing solution after every 30 minutes. The sensory epithelia are then mounted on the glass slide and 0.17mm cover glass using aqueous mounting media

**Table S3:** Primary and secondary antibodies used with the fixation condition.

| **Antibody** | **Source** | **Fixation** |
| --- | --- | --- |
| Myosin 7a | DSHB 138-1 | 4% PFA 1 hour RT or 4% PFA overnight |
| Arl13b | Proteintech 17711-1-AP | 4% PFA 1 hour RT or 4% PFA overnight |
| Beta Spectrin II | BD Biosciences 612562 | 4% PFA 1 hour RT or 4% PFA overnight |
| E Cadherin | BD Biosciences 610405 | 4% PFA 30minutes RT or Glyoxyl fixation 30minutes RT |
| N Cadherin | BD Biosciences 610920 | 4% PFA 30minutes RT or Glyoxyl fixation 30minutes RT |
| E Cadherin | Cell signalling Technology 3295 | 4% PFA 30minutes RT or Glyoxyl fixation 30minutes RT |
| P75NTR | Merck AB1554 | 4% PFA 1 hour RT |
| BLBP | Abcam ab32423 | 4% PFA 1 hour RT |
| Vangl2 | Sigma HPA027043 | 4% PFA 30minutes RT |
| p-RLC | Cell signalling Technology 3675 | 4% PFA 30minutes RT |
| pp-RLC | Cell signalling Technology 3674 | 4% PFA 30minutes RT and 2%TCA 30minutes on ice |
| Cdkn1b | sc-1641 | 4% PFA 1 hour RT |
| ZO1 | Sigma MABT339 | 4% PFA overnight or 4% PFA 30minutes RT and 2%TCA 30minutes on ice |
| Nectin 1 | MBL 48-12 | 4% PFA 30minutes RT |
| Nectin 2 | MBL 502-57 | 4% PFA 30minutes RT |
| R Cadherin | DSHB MRCD5 | 4% PFA 30minutes RT or Glyoxyl fixation 30minutes RT |
| E Cadherin Block | DSHB 7D6 | NA |
| N cadherin Block | DSHB 6B3 | NA |
| Phalloidin 488 | Invitrogen A12379 | 4% PFA 1 hour RT to 4% PFA overnight |
| Phalloidin 568 | Invitrogen A12380 | 4% PFA 1 hour RT to 4% PFA overnight |
| Goat anti-Rabbit 488 | Invitrogen A11034 | In all fixation conditions |
| Goat anti- Rabbit 594 | Invitrogen A11037 | In all fixation conditions |
| Goat anti-Rabbit 647 | Invitrogen A21244 | In all fixation conditions |
| Goat anti-mouse 488 | Invitrogen A-11001 | In all fixation conditions |
| Goat anti-mouse 594 | Invitrogen A-11005 | In all fixation conditions |
| Goat anti-mouse 647 | Invitrogen A-21235 | In all fixation conditions |
| Goat anti-rat 594 | Invitrogen A-11007 | In all fixation conditions |

### Imaging

The sensory epithelia were imaged using Olympus Fluoview 3000 inverted microscope controlled by Olympus FV31S-SW software at Central Imaging and Flow Cytometry Facility (CIFF) at NCBS. The images were obtained using 60X oil immersion objective of numerical aperture (NA) 1.42. The start and end point of the confocal volume were decided with the presence of stereocilia and junctions; images were obtained with a step size 0.5µm and pixel size as per Nyquist sampling criteria. The laser power was reduced using a neutral density filter (ND filter) to less than 10% and tuned from 0.1% to 10% for each experiment, keeping the voltage values between 400-500V, gain at 1 and offset between 2-3%, images were obtained by line sequential scanning. The light path was chosen such that the primary dichroic mirror and the secondary dichroic mirror were the same for each scanning and emitted light was collected by a high sensitivity spectral detector. For constructing the whole sensory epithelia images, the images were obtained using 10X air objective of NA 0.4 with step size of 1µm, line sequential scanning, HV values ranging between 400-500V.

The Olympus FV31S-SW software converts signals to 16-bit depth. The rotation feature of the software is used to align tissue wherever necessary.

### Ex-vivo organ culture

To culture cochlea, we used a three-dimensional collagen droplet culture previously explained (Singh et al 2022). Briefly, we made a collagen matrix solution by mixing 400µl of rat tail collagen I, 50µl of 10X DMEM, 30µl of 7.5% NaHCO3 and 5µl of HEPES using a pipette. Collagen mixture is poured as separate drops in 4-well dishes. The cochlea from the mouse embryo is dissected and a single cochlea is kept in each drop. Once all the drops have received one cochlea each the 4-well plate is transferred to a 37°C incubator maintaining 5% CO_2_ for 5 minutes. The plate is then taken out and the 500µl of culture media (1X DMEM supplemented with N2 and penicillin) is added using micropipettes and sterilised tips. The plate is then incubated at a 37°C incubator maintaining 5% CO_2_ for the required time. For small molecule perturbation, the solutions of the small molecule are added to the culture media of a well and treated as experiment and the solvent of the compound are added to the culture media and treated as control. For blocking experiments, Bovine Serum Albumin (BSA) was used as a control. The concentration of small molecule inhibitors and blocking antibodies is presented in table S4.

**Table S4:** Concentration of small molecule inhibitors used for ex-vivo organ culture.

| Small Molecule Inhibitor | Identifier | Concentration used |
| --- | --- | --- |
| ML7 | Sigma 475880 | 25μM |
| E cadherin | DSHB 7D6 | 5μl/ml (10μg/ml) |
| N cadherin Block | DSHB 6B3 | 5μl/ml (10μg/ml) |
| Su5402 | Calbiochem 572630 | 25μM |

### Image Analysis

The confocal images obtained were opened in FIJI (NIH Image J) and processed to form single-channel and multichannel merged images. Images were provided with a scale bar using the metadata confocal file. To evaluate the morphological features of the cells in the epithelia we segmented the images using the Tissue-Analyzer plugin and used the segmented images to obtain values of **apical surface area, shape index, circularity, Feret angle, aspect ratio** using the auto-measure tool in Image J. We also obtained other morphological features of the epithelia

A. **Polarity:** The position of kinocilia is ascertained by the Beta-spectrin2 or Arl13b staining. The X, Y coordinates of the cilia position is determined using Image J and a vector is drawn from the centroid of HCs to the cilia position. This vector length (r) is normalised to the radius of cells and the angle (θ) is with respect to the P-D axis (θ). The cilia position is then plotted along with the calculation of the mean angle (denoted by the angle of the arrow) and circular standard deviation (denoted by the length of the arrow) using the custom-made script, which would be deposited on GitHub after the acceptance of this manuscript.
B. **4-Cell vertex calculations:** The segmented images in Tissue analyser are processed to obtain the data for bonds using SQL command. The data for bond position and vertex coordinates are tabulated in MS Excel. Using a MS Excel macro, the vertex with more than 4 cells and the total number of vertices are computed.
C. **Width of the compartments:** For calculation of sensory compartment width a straight line is drawn in FIHI from the medial edge of IHC to the lateral edge of border cells. The medial sensory compartment is calculated from medial edge of IHC to lateral edge of pillar cells; the lateral sensory compartment is calculated from medial edge of OHC1 to lateral edge of border cells.
D. **Tortuosity.** Using the centroid of IHC a line was drawn connecting 10 IHC and its length (L) was calculated and the shortest distance between the IHC was calculated between the first and last IHC (l). The tortuosity was calculated as ratio of L/l.

### Visualisation

Graphs are made using Prism-GraphPad. Rose stack plots representing the axis of elongation for KO cells are plotted using Oriana software from Kovach computing services. The images are assembled into figures using Inkscape.

### Statistics

We performed a two-tail unpaired T-test without assuming Gaussian distribution as the Mann-Whitney T-test to calculate the significance level using Prism-GraphPad software.

## ACKNOWLEDGEMENTS

This work was supported by the Department of Atomic Energy, Government of India, Project Identification No. RTI 4006, and grants from SERB, TIFR Infosys-Leading Edge Grant, the Royal National Institute for Deaf People, and the International Foundation for Research and Education, via a Simons-Ashoka ECF fellowship to AP, for this research. We thank Central Imaging and Flowcytometry Facility (CIFF), Animal Care and Resource Centre (ACRC) and laboratory kitchen at NCBS. We thank Srikala Raghvan for providing the Vinculin mouse. We thank Hiroshi Hamada and Sandeep Krishna for comments on the manuscript. A. P. thanks ICTP - ICTS Winter School on Quantitative Systems Biology for introducing quantitative morphogenesis. The Myo7a antibody developed by Dana J. Orten, E-cadherin antibody developed by Warren Gallin, and the N-cadherin antibody developed by Karen A Knudsen was obtained from the Developmental Studies Hybridoma Bank, created by the NICHD of the NIH and maintained at The University of Iowa, Department of Biology, Iowa City, IA 52242. Cdkn1b antibody was a kind gift from Dr Dimple Notani. We are grateful to Sweety Meel and the members of Ear Lab for their feedback.

## AUTHOR CONTRIBUTIONS

A.P. and R.K.L conceived the project and designed the experiments. A.P performed experiments and analysed the data. A.P. and S.R. performed Vangl2 lgr5 experiments. S.R. and R.K helped in generating different mouse strains and provided inputs in experimental design and manuscript preparation. P.M performed EdU experiments. A.S.I and A.P. developed codes to analyse data. A.P. and R.K.L wrote and revised the manuscript.

## Supplementary Figures

**Figure S1.**
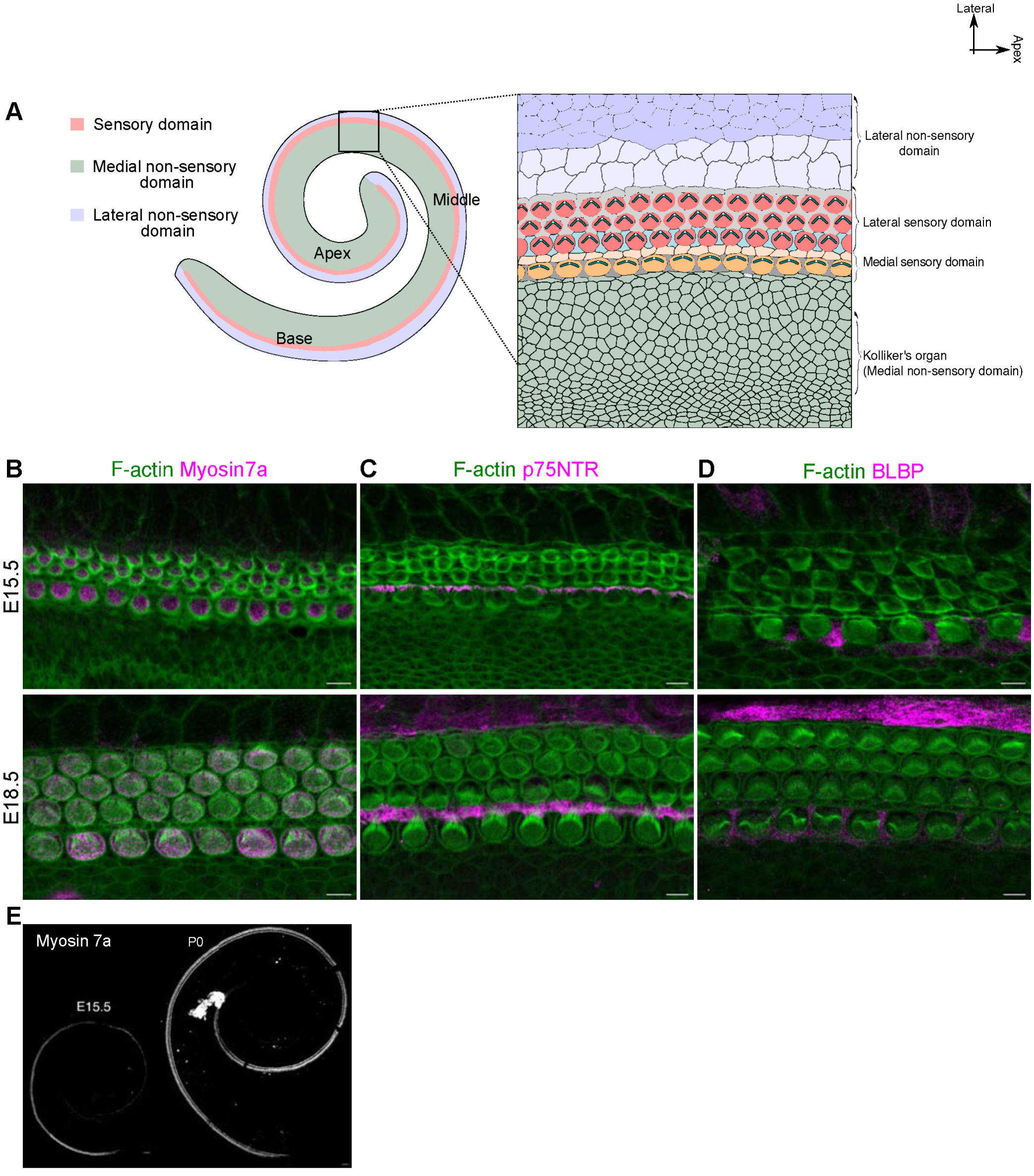
Radial patterning of sensory and non-sensory domains is preserved during cochlear elongation. A. Schematic representing the organization of various sensory and non-sensory domains and their constituent cell types in the mouse organ of Corti. B. Base of E15.5 and E18.5 OC stained for F-actin (green) and Myosin7a (magenta). C. Base of E15.5 and E18.5 OC stained for F-actin (green) and p75NTR (magenta). D. Base of E15.5 and E18.5 OC stained for F-actin (green) and BLBP (magenta). E. OC from E15.5 and PO stained for Myosin 7a. Scale Bar: 50µm in E and 5µm in B-D Image orientation: Top is lateral, Right is Apex.

**Figure S2.**
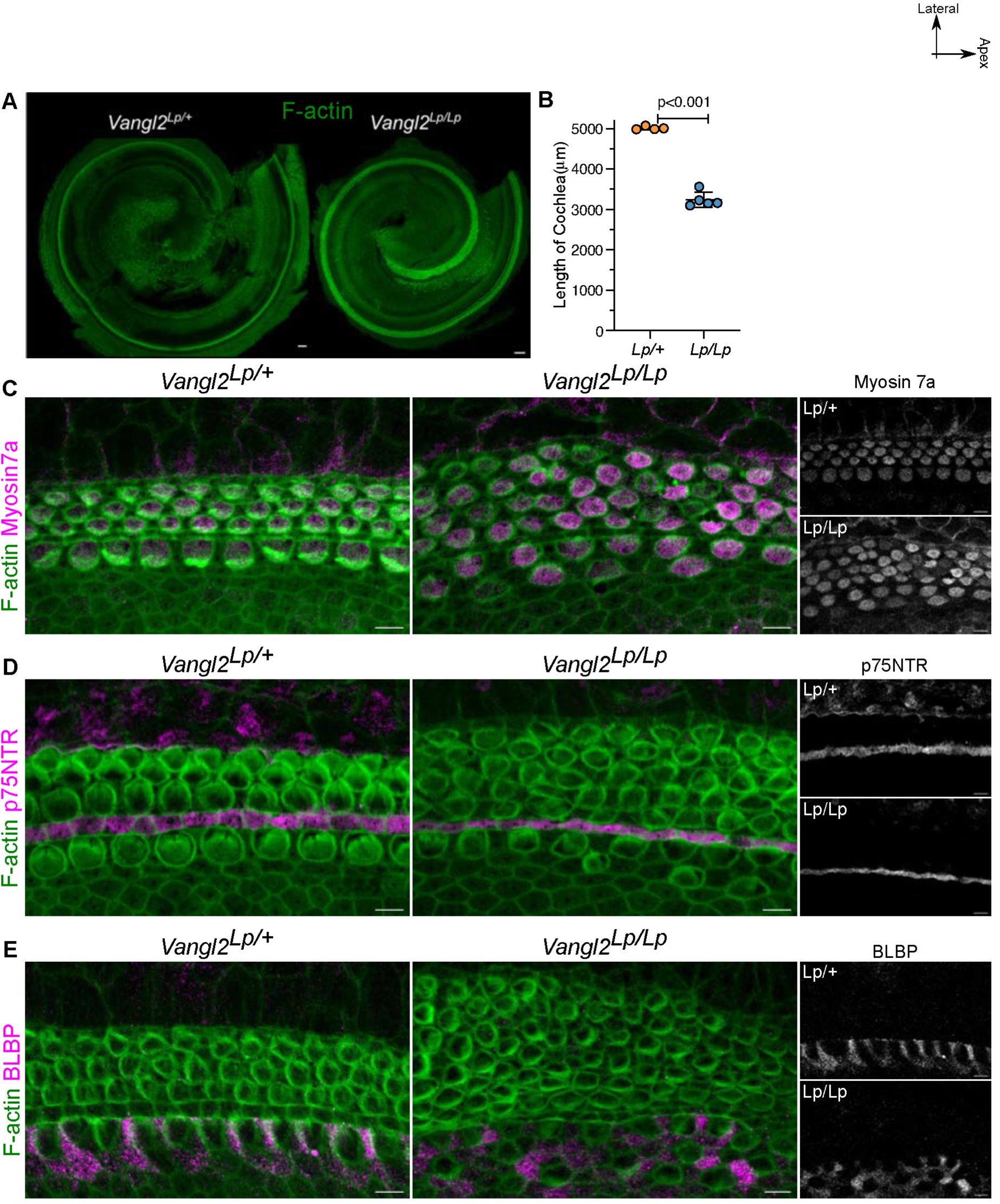
Domain organisation is maintained even at the apical turn in looptail mutants. A. OC from embryonic day (E)18.5 heterozygous (*Vangl2 ^Lp/+^*) and homozygous (*Vangl2 ^Lp/Lp^*) looptail mutant stained for f-actin (green). B. Length of E18.5 cochlea from heterozygous (*Vangl2 ^Lp/+^*) and homozygous (*Vangl2 ^Lp/Lp^*) looptail mutant. N=4 / 5 (Het/Homo) C. Apex of E18.5 OC from heterozygous (*Vangl2 ^Lp/+^*) and homozygous (*Vangl2 ^Lp/Lp^*) looptail mutant stained for F-actin (green) and Myosin 7a (magenta and grey). N= 4. D. Apex of E18.5 OC from heterozygous (*Vangl2 ^Lp/+^*) and homozygous (*Vangl2 ^Lp/Lp^*) looptail mutant stained for F-actin (green) and p75NTR (magenta and grey). N= 4. E. Apex of E18.5 OC from heterozygous (*Vangl2 ^Lp/+^*) and homozygous (*Vangl2 ^Lp/Lp^*) looptail mutant stained for F-actin (green) and BLBP (magenta and grey). N= 4. Scale Bar: 50µm in B and 5µm in C-E Unpaired T-test Image orientation: Top is lateral, Right is Apex.

**Figure S3.**
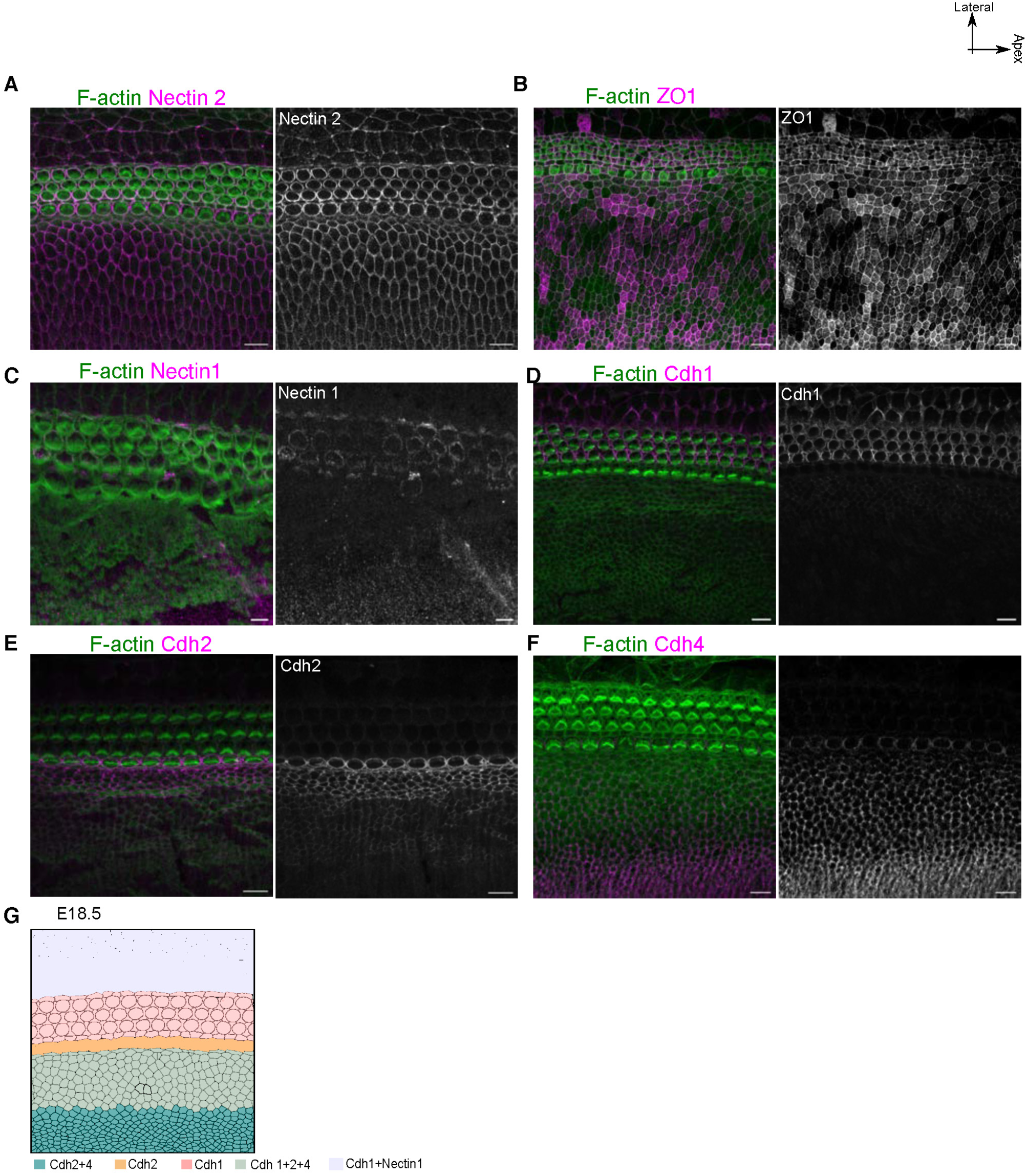
Differential expression of adhesion molecule super-impose on domain identity. A. E18.5 OC stained for F-actin (green) and Nectin2 (magenta and grey). B. E18.5 OC stained for F-actin (green) and Zonula Occludens-1 (ZO1)(magenta and grey). C. E18.5 OC stained for F-actin (green) and Nectin2 (magenta and grey). D. E18.5 OC stained for F-actin (green) and Cdh1 (magenta and grey). E. E18.5 OC stained for F-actin (green) and Cdh2 (magenta and grey). F. E18.5 OC stained for F-actin (green) and Cdh4 (magenta and grey). G. Schematic representing the combinatorial expression of adhesion molecule super-imposed on the cell types of OC. Scale Bar: 10µm Image orientation: Top is lateral, Right is Apex.

**Figure S4.**
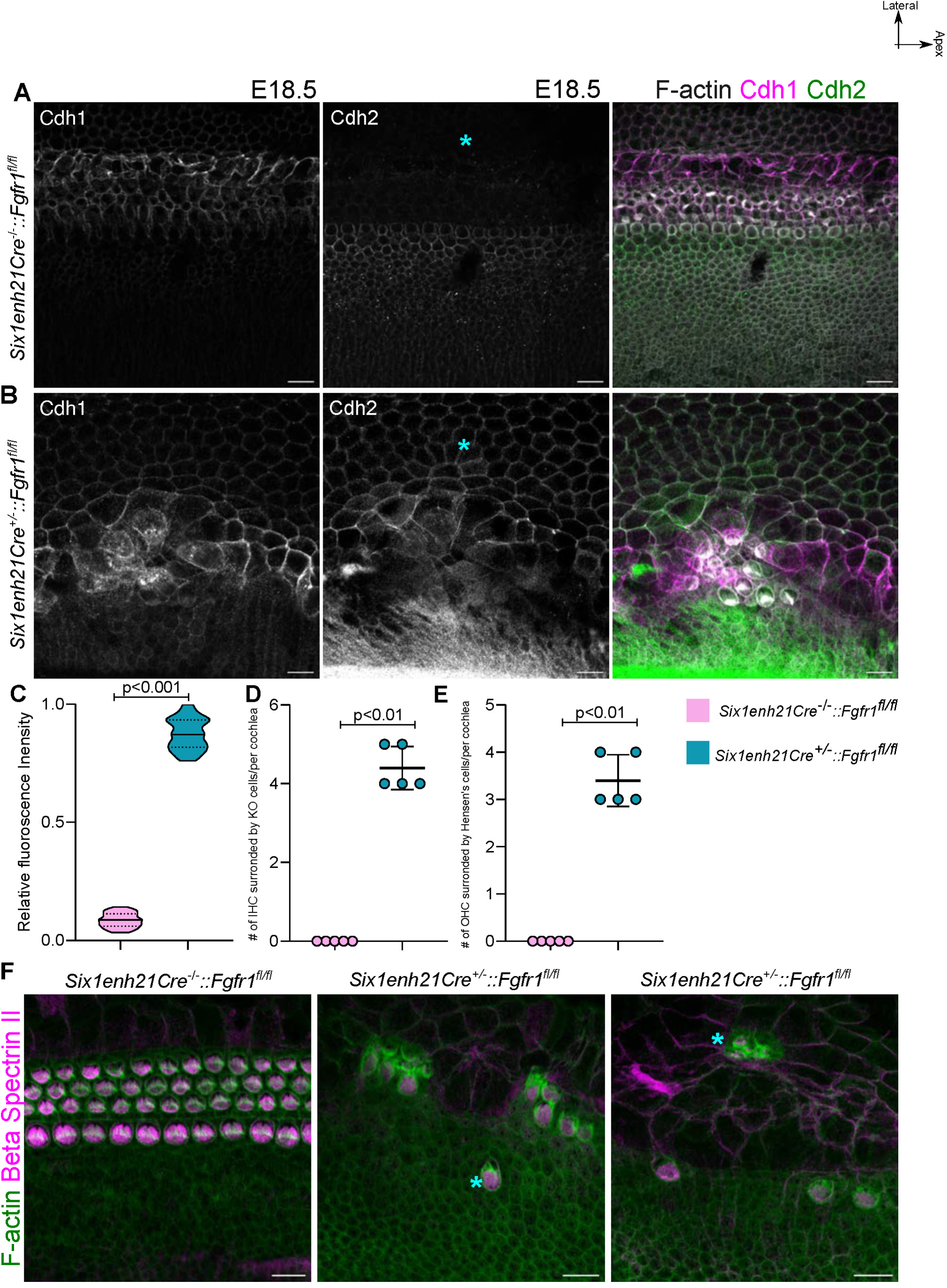
Disruption of Fgfr1 signalling leads to sustained misexpression of cadherin and disruption in organisation. A. E18.5 OC from control embryos (*Six1enh21Cre^−/−^::Fgfr1^fl/fl^)* stained for Cdh1 (grey and magenta), Cdh2 (grey and green) and F-actin (grey in merged). Asterisk indicates the absence of Cdh2 signals from lateral non-sensory domain. N=4 embryos B. E18.5 OC from Fgfr1 mutant embryos (*Six1enh21Cre^+/−^::Fgfr1^fl/fl^)* stained for Cdh1 (grey and magenta), Cdh2 (grey and green) and F-actin (grey in merged). Asterisk indicates the ectopic Cdh2 signals from lateral non-sensory domain. N=4 embryos C. Relative Fluorescence Intensity of Cdh2 in Claudius cells in control and Fgfr1 mutant cochlea at E18.5. D. Number of IHC surrounded by KO cells on all side per cochlea in control and fgfr1 mutant cochlea. This shows medial non-sensory domain is intermixed with the medial sensory domain. E. Number of OHC surrounded by Hensen’s cells on all side per cochlea in control and fgfr1 mutant cochlea. This shows lateral non-sensory domain is intermixed with the lateral sensory domain. F. E18.5 OC from control and Fgfr1 mutant embryos (*Six1enh21Cre^+/−^::Fgfr1^fl/fl^)* stained for F-actin (green) and HC marker Beta Spectrin II (magenta). Asterisk indicates the presence of HCs in the lateral and medial non-sensory domains. N=5 embryos Unpaired T-test Scale Bar: 10µm Image orientation: Top is lateral, Right is Apex.

**Figure S5.**
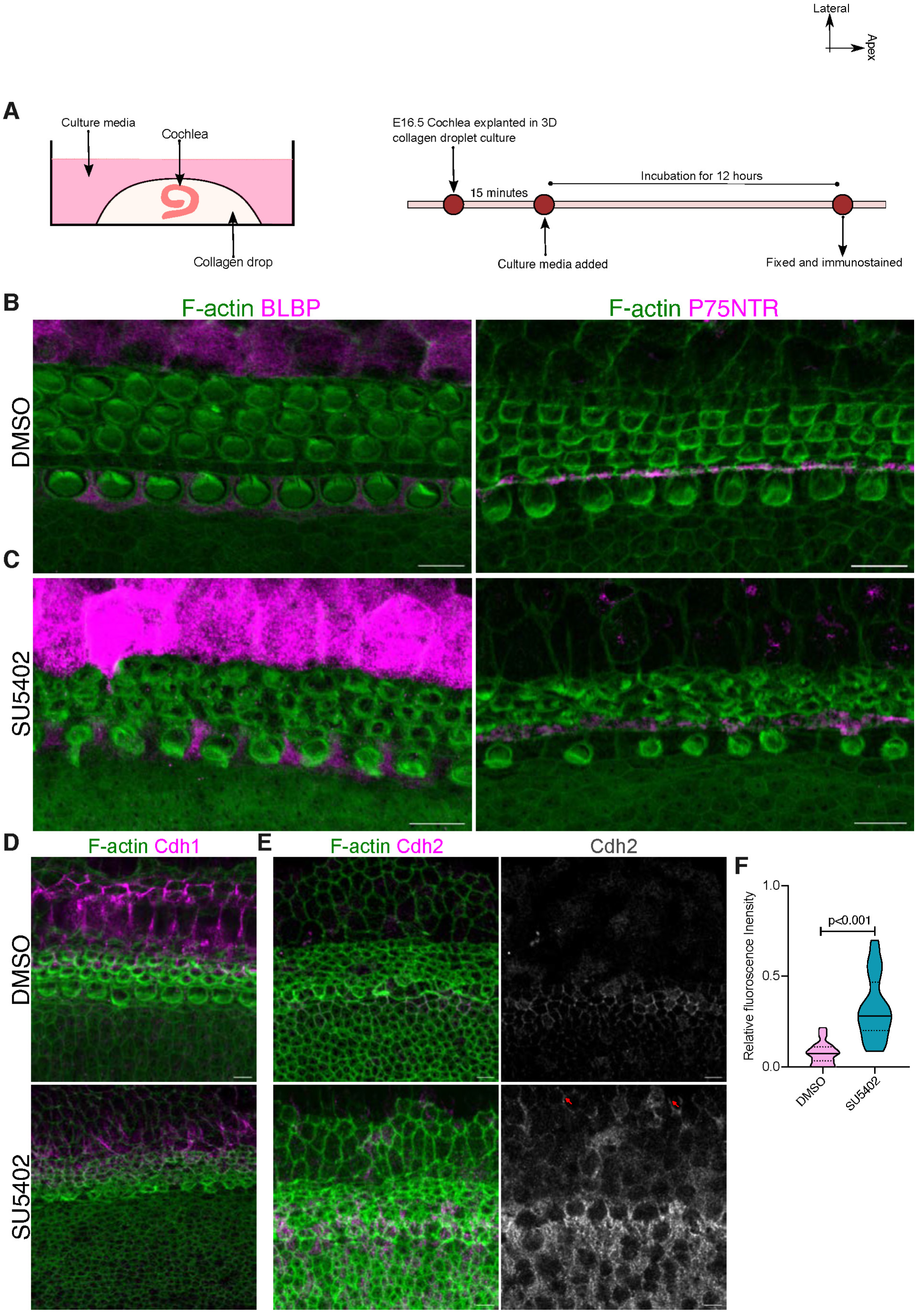
Disruption of FGF signalling drives misexpression of Cdh2 in lateral non-sensory domain. A. Schematic representing the collagen droplet culture and treatment condition for an ex-vivo explant culture of OC. B. E16.5 Cochlea cultured in presence of DMSO for 12 hours stained for F-actin (green), BLBP (magenta) and p75NTR (magenta). C. E16.5 Cochlea cultured in presence of Fgfr1 inhibitor (Su5402, 25µM) stained for F-actin (green), BLBP (magenta) and p75NTR (magenta). D. E16.5 Cochlea cultured in presence of Fgfr1 inhibitor (Su5402, 25µM) stained for F-actin (green), Cdh1(magenta). E. E16.5 Cochlea cultured in presence of Fgfr1 inhibitor (Su5402, 25µM) stained for F-actin (green), Cdh2 (magenta, grey) showing misexpression of Cdh2 in Claudius cells similar to the genetic perturbation in Fig S4B. F. Relative Fluorescence Intensity of Cdh2 in Claudius cells in control (DMSO treated) and Fgfr1 inhibited (Su5402 treated) cochlea at E18.5. Unpaired T-test Scale Bar: 10µm Image orientation: Top is lateral, Right is Apex.

**Figure S6.**
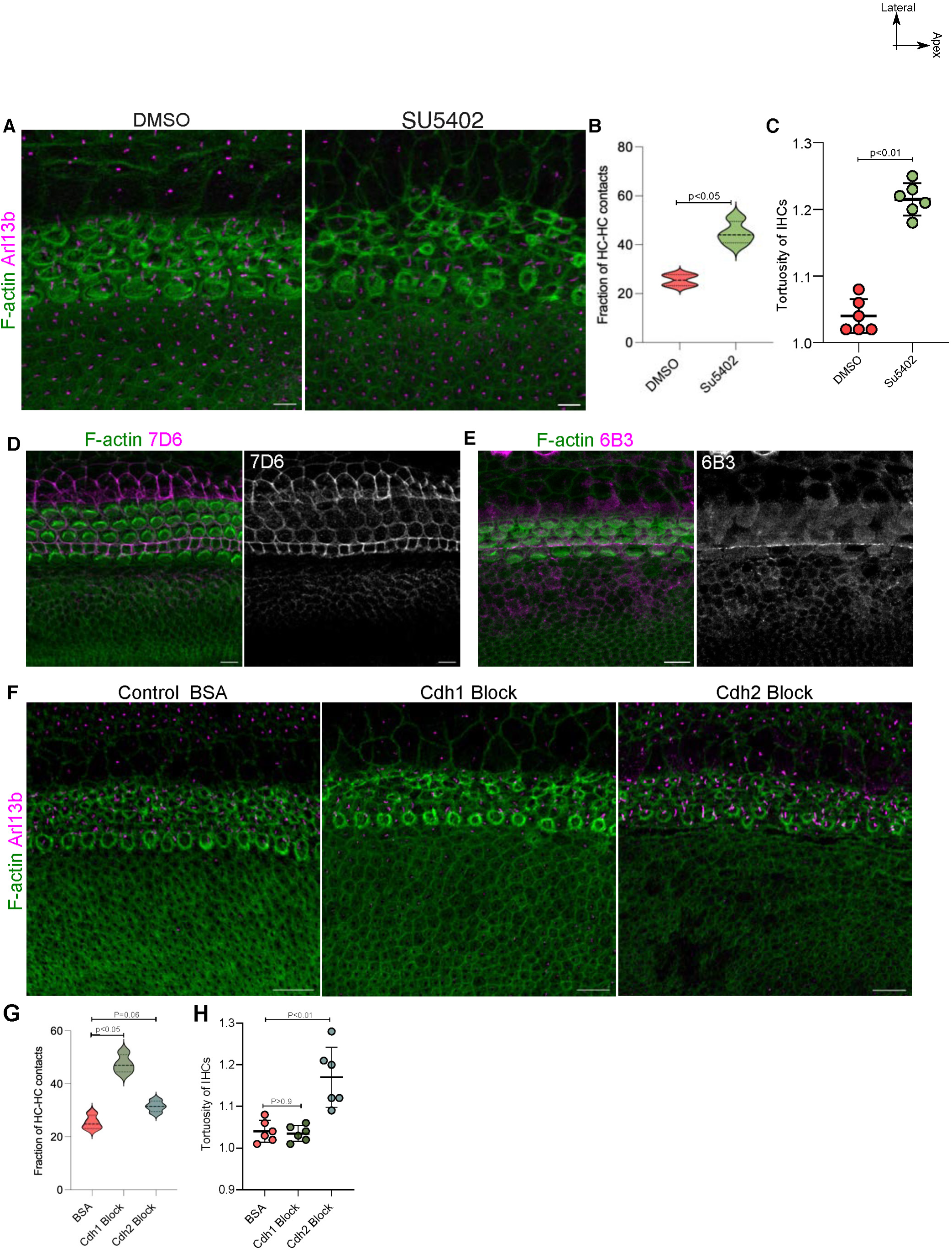
FGF inhibited and cadherin blocked OC showed disorganisation of domains. A. E16.5 Cochlea cultured in presence of DMSO and Su5402 for 12 hours stained for F-actin (green), BLBP (magenta). B. Fraction of HC-HC contacts in DMSO treated and Su5402 treated OC. C. Straightness of IHC represented by low tortuosity in DMSO treated cochlea and the disruption of this straightness represented by high tortuosity in Su5402 treated cochlea. D. Cochlea cultured for 1 hour in presence of cadherin blocking antibodies 7D6, which block interactions among Cdh1, stained using F-actin (green) and secondary antibodies (magenta, grey). E. Cochlea cultured for 1 hour in presence of cadherin blocking antibodies 6B3, which block interactions among Cdh2, stained using F-actin (green) and secondary antibodies (magenta, grey). F. 12-hour explant of E15.5 OC in presence of Bovine Serum Albumin (BSA, 0.1%), Cdh1 blocking antibodies (7D6, 10ug/ml), Cdh2 blocking antibodies (6B3, 10ug/ml) stained for F-actin (green) and Arl13b (magenta). N=4 cochlea G. Fraction of HC-HC contacts in BSA treated, Cdh1 blocked and Cdh2 blocked OC. H. Tortuosity of IHC in BSA treated, Cdh1 blocked and Cdh2 blocked OC. Unpaired T-test Scale Bar: 10µm and Image orientation: Top is lateral, Right is Apex.

**Figure S7.**
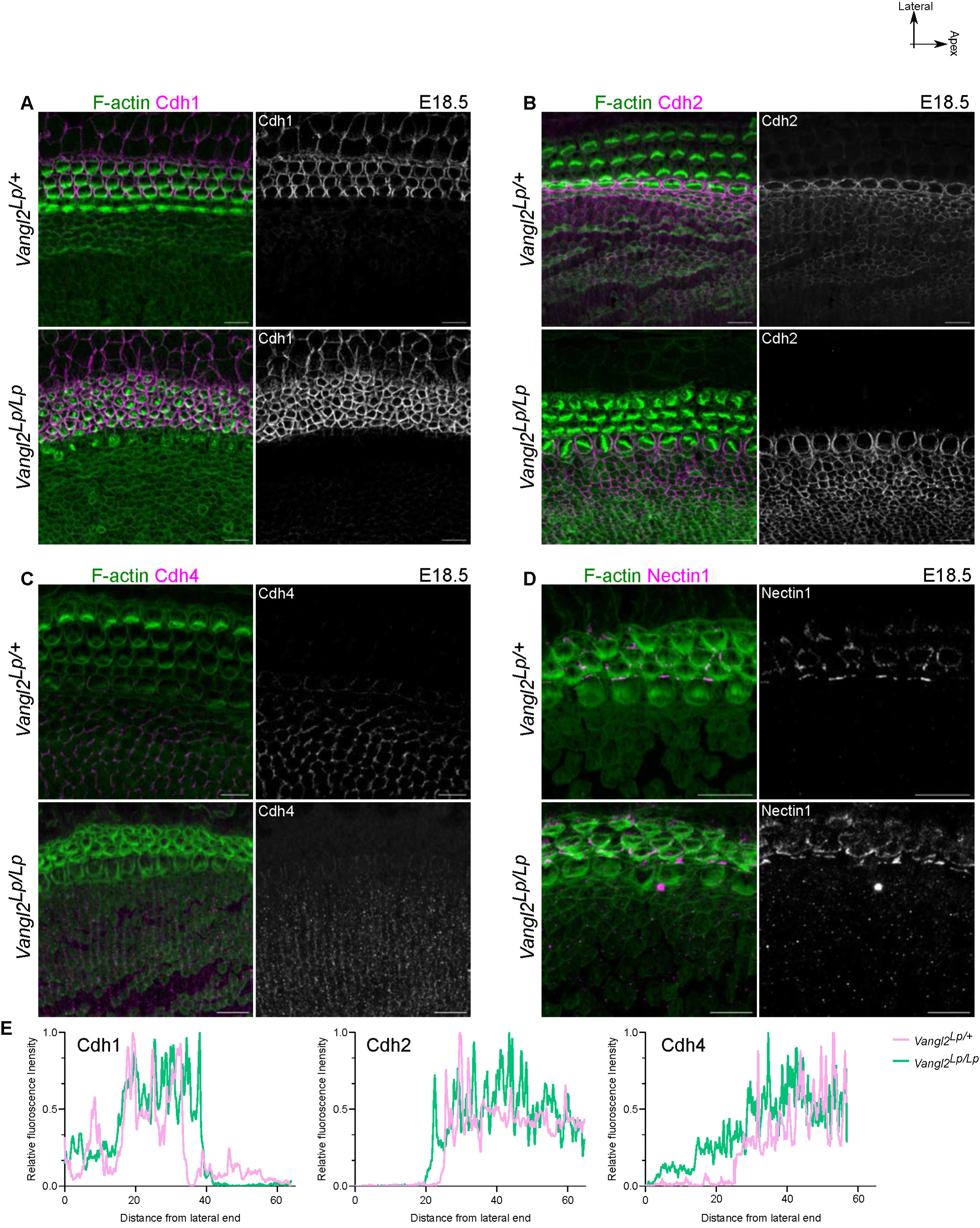
Adhesion code is maintained in looptail mutants with defective convergent extension. A. E18.5 OC from heterozygous (*Vangl2 ^Lp/+^*) and homozygous (*Vangl2 ^Lp/Lp^*) looptail mutant stained for F-actin (green) and Cdh1 (magenta and grey). N= 3 cochlea B. E18.5 OC from heterozygous (*Vangl2 ^Lp/+^*) and homozygous (*Vangl2 ^Lp/Lp^*) looptail mutant stained for F-actin (green) and Cdh2 (magenta and grey). N= 3 cochlea C. E18.5 OC from heterozygous (*Vangl2 ^Lp/+^*) and homozygous (*Vangl2 ^Lp/Lp^*) looptail mutant stained for F-actin (green) and Cdh4 (magenta and grey). N= 3 cochlea D. E18.5 OC from heterozygous (*Vangl2 ^Lp/+^*) and homozygous (*Vangl2 ^Lp/Lp^*) looptail mutant stained for F-actin (green) and Nectin1 (magenta and grey). N= 3 cochlea E. Relative fluorescence intensity of Cdh1, Cdh2 and Cdh4 along the medio-lateral axis of OC at E18.5 from heterozygous (*Vangl2 ^Lp/+^*) and homozygous (*Vangl2 ^Lp/Lp^*) looptail mutant in pink and green, respectively. Scale Bar: 10µm Image orientation: Top is lateral, Right is Apex.

**Figure S8.**
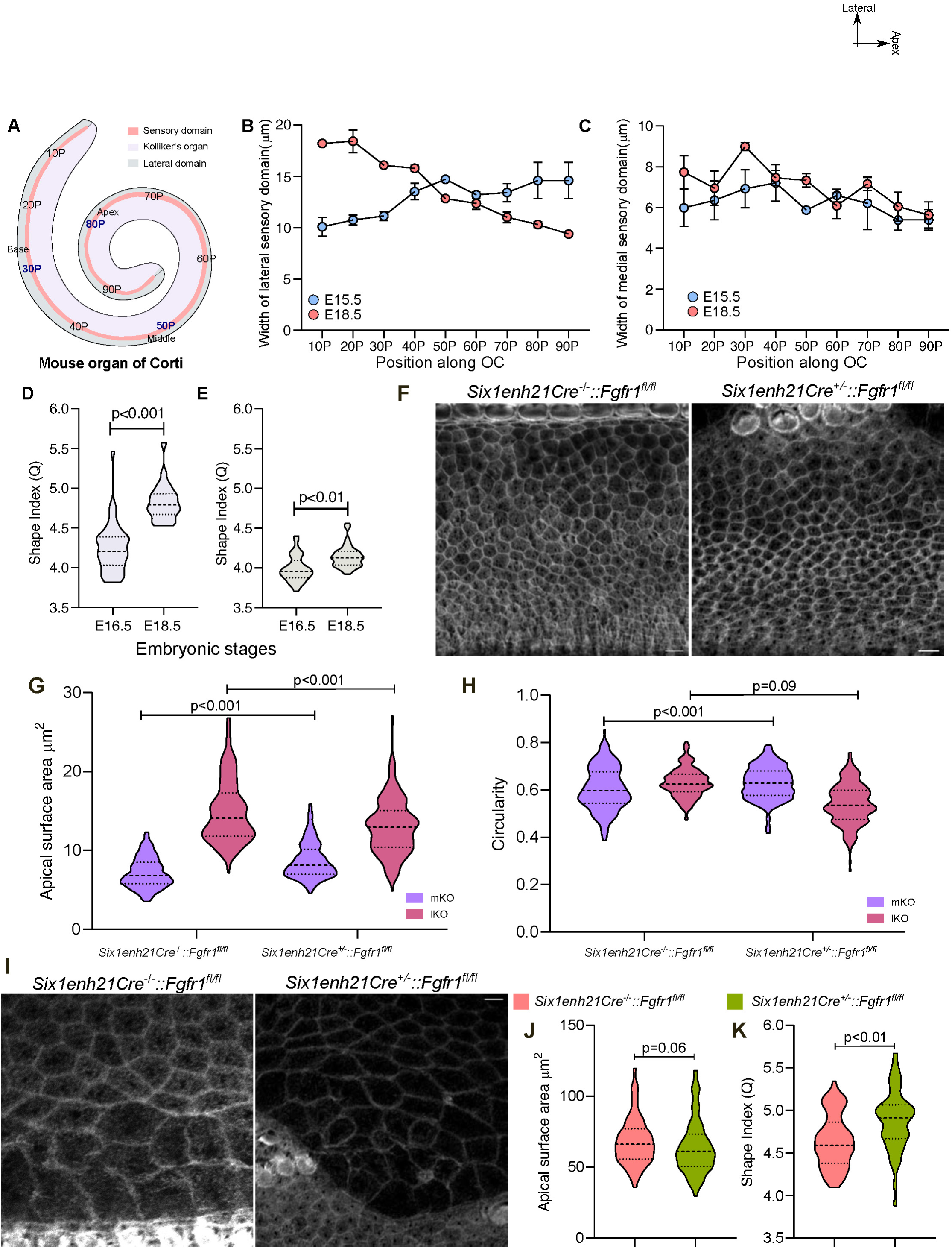
Adhesion code ensures discrete organisation of each compartments. A. Schematic of OC representing the 9 equidistant points along the base-apex axis of the OC. B. Width of the lateral sensory domain along the 9 positions along the base-apex axis at E15.5 and E18.5. N=4 cochlea each stage C. Width of the medial sensory domain along the 9 positions along the base-apex axis at E15.5 and E18.5. N=4 cochlea each stage D. Shape index (Q=perimeter/sqrt of area) of Hensen’s Cells at E16.5 and E18.5. N=150/169 for Hensen and 144/153 for Claudius (E16.5/E18.5). E. Shape index (Q=perimeter/sqrt of area) of Claudius Cells at E16.5 and E18.5. N= 144/153 (E16.5/E18.5). F. E18.5 base of OC from control (*Six1enh21Cre^−/−^::Fgfr1^fl/fl^)* and Fgfr1 mutant embryos (*Six1enh21Cre^+/−^::Fgfr1^fl/fl^)* stained for F-actin (grey) showing KO domain. N=4 G. Apical surface area of mKO and lKO cells from control (*Six1enh21Cre^−/−^::Fgfr1^fl/fl^)* and Fgfr1 mutant embryos (*Six1enh21Cre^+/−^::Fgfr1^fl/fl^).* N=150/216 for control and 209/262 for mutant (mKO/lKO). H. Circularity of mKO and lKO cells from control (*Six1enh21Cre^−/−^::Fgfr1^fl/fl^)* and Fgfr1 mutant embryos (*Six1enh21Cre^+/−^::Fgfr1^fl/fl^).* N=150/216 for control and 209/262 for mutant (mKO/lKO). I. E18.5 base of OC from control (*Six1enh21Cre^−/−^::Fgfr1^fl/fl^)* and Fgfr1 mutant embryos (*Six1enh21Cre^+/−^::Fgfr1^fl/fl^)* stained for F-actin (grey) showing lateral non-sensory domain. N=4 J. Apical surface area of Hensen’s Cells from control (*Six1enh21Cre^−/−^::Fgfr1^fl/fl^)* and Fgfr1 mutant embryos (*Six1enh21Cre^+/−^::Fgfr1^fl/fl^).* N=65/98. (control/mutant). K. Shape index of Hensen’s Cells from control (*Six1enh21Cre^−/−^::Fgfr1^fl/fl^)* and Fgfr1 mutant embryos (*Six1enh21Cre^+/−^::Fgfr1^fl/fl^).* N=65/98. (control/mutant). Scale Bar: 10µm Unpaired T-test Image orientation: Top is lateral, Right is Apex. *Six1enh21Cre^−/−^* means cre negative and *Six1enh21Cre^+/−^* cre positive

**Figure S9.**
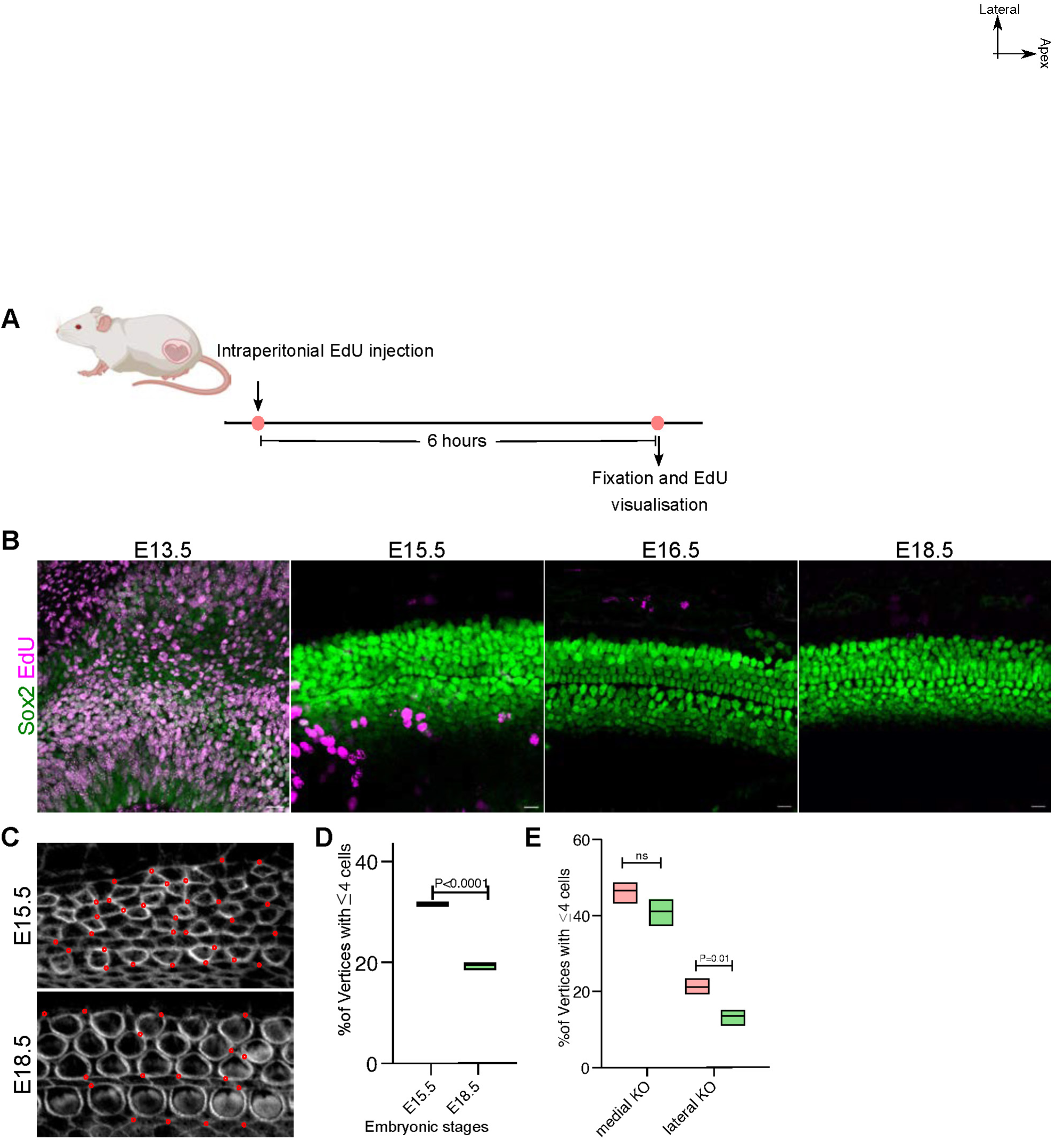
Cellular intercalation drives compartment specific reorganisation. A. Schematic representing the timeline of EdU injection and staining. B. OC of embryos from E13.5, E15.5, E16.5 and E18.5 pregnant females injected with EdU stained for Sox2 (green) to mark sensory epithelia and click-chemistry based EdU (Magenta). N=3 C. OC from E15.5 and E18.5 base position (30P) stained for F-actin (grey), overlayed with red circles representing vertex with 4 or more cells. D. Percentage of vertex with 4 or more cells in the sensory domain at E15.5 and E18.5. N=3 embryos each with 205/615 for E15.5 and 122/625 for E18.5 (4 or more cell vertices /Total vertices). E. Percentage of vertex with 4 or more cells in the medial and lateral KO domain at E15.5 and E18.5. N= 3 cochlea each, 498/1070 mKO at E15.5 and 259/630 at E18.5; 318/1532 lKO at E15.5 and 111/835 at E18.5 (4 or more cell vertices /Total vertices) Scale Bar: 20µm for E13.5 and 10µm for rest. Unpaired T-test, ns= P>0.05 Image orientation: Top is lateral, Right is Apex.

**Figure S10.**
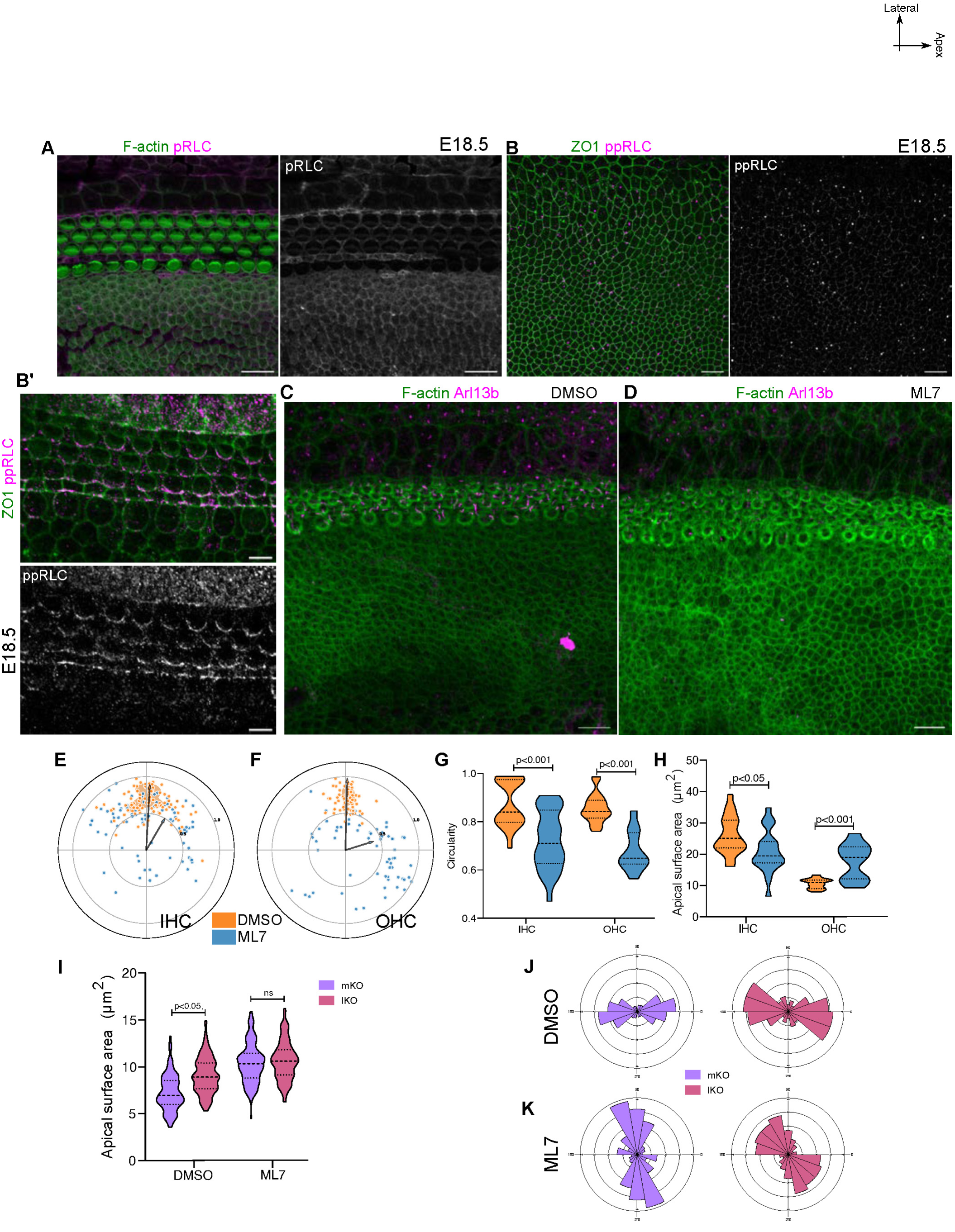
NMII-activity drives organisation within each compartment. A. E18.5 OC stained for F-actin (green) and mono-phosphorylated form of RLC (pRLC, magenta, grey). B. KO and Sensory domain from E18.5 OC stained for ZO1(green) and di-phosphorylated form of RLC (ppRLC, magenta, grey). C. E16.5 cochlea cultured ex-vivo for 8 hours in a 3D-collagen droplet culture with DMEM supplemented with DMSO, stained for F-actin (green) and Arl13b (magenta). N=6 cochlea D. E16.5 cochlea cultured ex-vivo for 8 hours in a 3D-collagen droplet culture with DMEM supplemented with or 25µM ML7, stained for F-actin (green) and Arl13b (magenta). N=6 cochlea E. Polar coordinates representing position of kinocilia of IHC from OC cultured in presence (blue) or absence (orange) of MLCK-inhibitor ML7. N= 66/68 (DMSO/ML7) F. Polar coordinates representing position of kinocilia of OHC from OC cultured in presence (blue) or absence (orange) of MLCK-inhibitor ML7. N= 68/61 (DMSO/ML7) G. Circularity of IHC and OHC from OC cultured in presence (blue) or absence (orange) of MLCK-inhibitor ML7. N=58/61 for IHC and 69/66 for OHC (DMSO/ML7) H. Apical surface area of IHC and OHC from OC cultured in presence (blue) or absence (orange) of MLCK-inhibitor ML7. N=58/61 for IHC and 69/66 for OHC (DMSO/ML7). I. Apical surface area of mKO and lKO cells from OC cultured in presence or absence of MLCK-inhibitor ML7. N= 193/207 for DMSO and 203/214 for ML7 (DMSO/ML7). J. Rose stack plot representing the axis of cell elongation for mKO and lKO cells in OC cultured in DMSO. N= 193/207 mKO/lKO. K. Rose stack plot representing the axis of cell elongation for mKO and lKO cells in OC cultured in ML7. N= 203/214 mKO/lKO. Image orientation: Top is lateral, Right is Apex. Scale Bar: 10µm Unpaired T-test, ns=non-signficant, P>0.05

**Figure S11.**
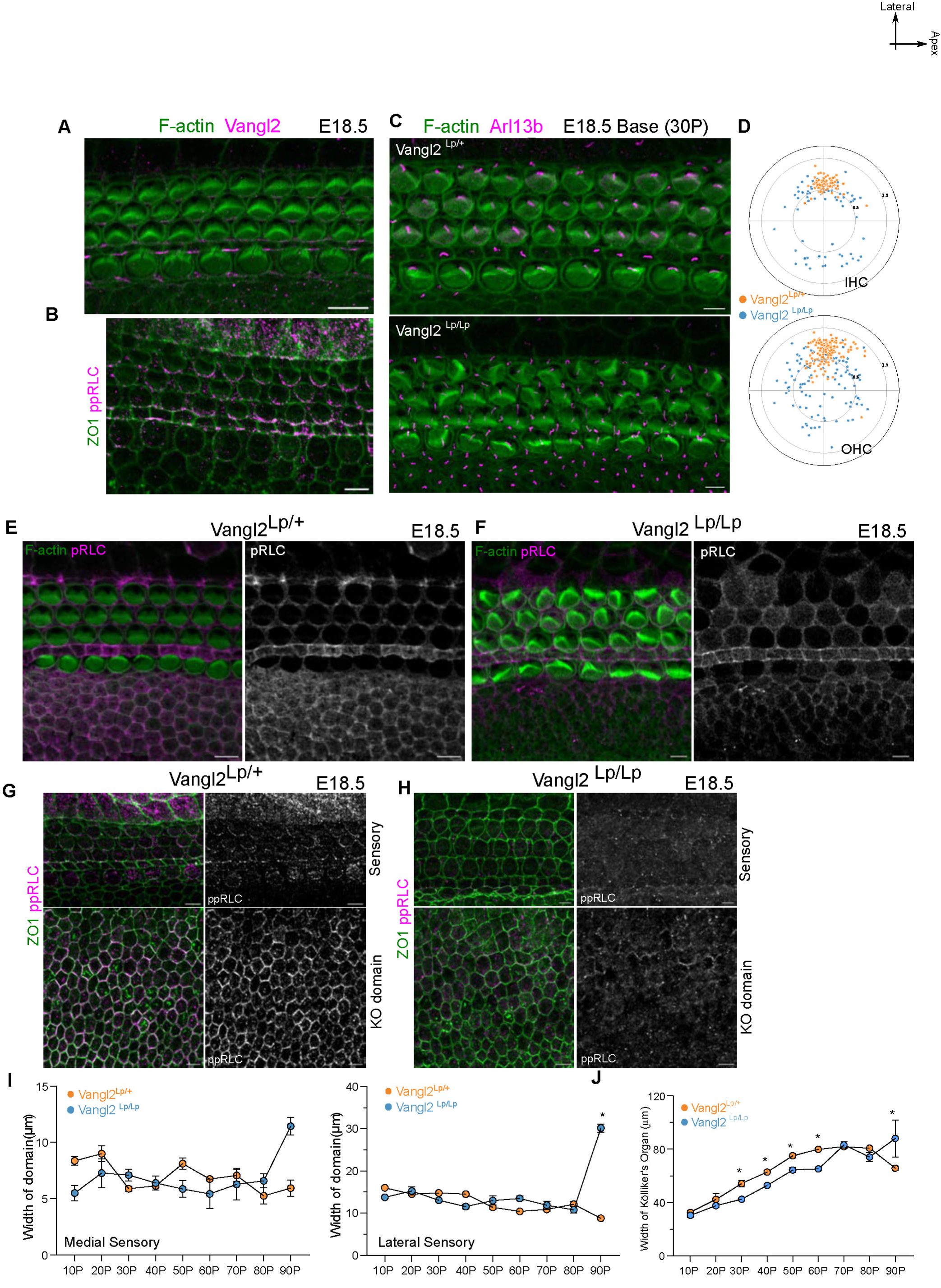
Vangl2 driven NMII-activity drives domain scale and cellular scale organisation. A. Sensory domain from E18.5 OC stained for F-actin (green) and Vangl2 (magenta). B. Sensory domain from E18.5 OC stained for ZO1(green) and di-phosphorylated form of RLC (ppRLC, magenta, grey). C. Base region (30P) of E18.5 OC from heterozygous (*Vangl2 ^Lp/+^*) and homozygous (*Vangl2 ^Lp/Lp^*) looptail mutant stained for F-actin (green) and Arl13b (magenta). N= 6 cochlea. D. Polar coordinates of kinocilia of IHC and OHC at Base region (30P) of E18.5 OC from heterozygous (*Vangl2 ^Lp/+^*) in orange and homozygous (*Vangl2 ^Lp/Lp^*) looptail mutant in blue. N=66/66 for IHC and N=110/110 for OHC (Het/Homo). E. E18.5 OC from heterozygous looptail mutant stained for F-actin (green) and pRLC (magenta, grey). F. E18.5 OC from homozygous looptail mutant stained for F-actin (green) and pRLC (magenta, grey). G. E18.5 OC from heterozygous looptail mutant stained for ZO1(green) and ppRLC (magenta, grey). H. E18.5 OC from homozygous looptail mutant stained for ZO1(green) and ppRLC (magenta, grey). I. Width of medial and lateral sensory domain along the OC from heterozygous (*Vangl2^Lp/+^*) and homozygous (*Vangl2 ^Lp/Lp^*) looptail mutant at E18.5. N=4 embryos J. Width of Kölliker’s organ along the OC from heterozygous (*Vangl2 ^Lp/+^*) and homozygous (*Vangl2 ^Lp/Lp^*) looptail mutant at E18.5. N=4 embryos. Image orientation: Top is lateral, Right is Apex. Scale Bar: 10µm Unpaired T-test, ns=non-signficant, P>0.05, *= P<0.05

**Figure S12.**
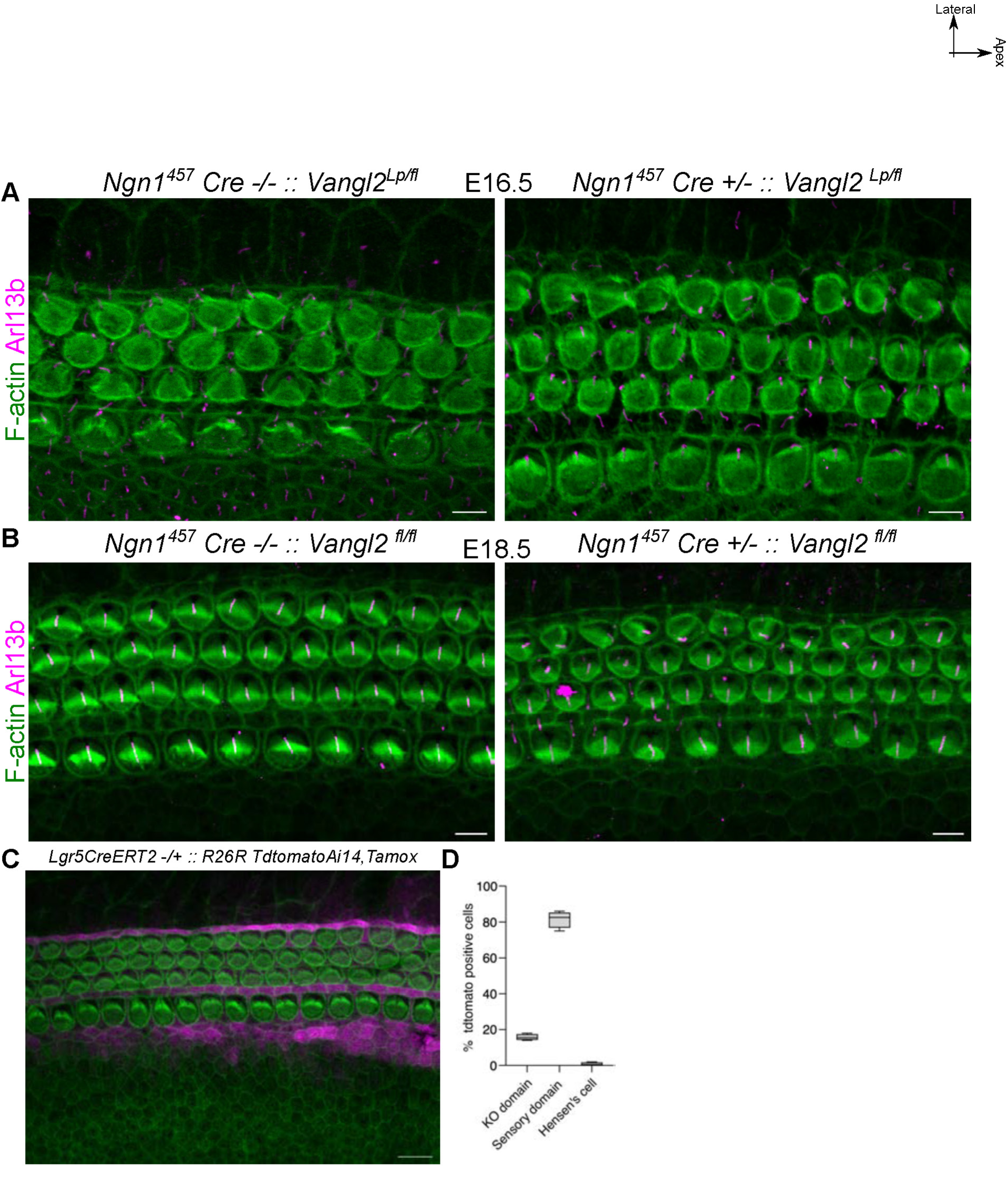
Compartment specific deletion of Vangl2 has non-linear effects on other compartments. A. Base region (30P) of E16.5 OC from Cre negative control (*Ngn1^457^-Cre^−/−^:: Vangl2^lp/fl^*) and Ngn1^457^cre positive mutant (*Ngn1^457^-Cre^+/−^:: Vangl2^lp/fl^*) stained for F-actin (green) and Arl13b (magenta). B. Base region (30P) of E18.5 OC from Cre negative control (*Ngn1^457^-Cre^−/−^:: Vangl2^fl/fl^*) and Ngn1^457^cre positive mutant (*Ngn1^457^-Cre^+/−^:: Vangl2^fl/fl^*) stained for F-actin (green) and Arl13b (magenta). Note: here both copies of Vangl2 is flox allele. C. E18.5 OC from Lgr5CreERT2::Ai14, induced with Tamoxifen stained with F-actin (green) showing cre mediated expression of Tdtomato (magenta). D. Percentage of tdtomato positive cells from the Ngn1^457^-Cre::Ai14 cochlea in KO, sensory and lateral non-sensory domain. N=3 Scale Bar: 10µm Image orientation: Top is lateral, Right is Apex.

